# Intron-mediated enhancement boosts Rtn4 circRNA expression: A robust method for exploring circRNA function

**DOI:** 10.1101/257105

**Authors:** Dingding Mo, Xinping Li, Di Cui, Jeanne-Franca Vollmar

## Abstract

CircRNAs are expressed in many important biological processes. Studying their function requires an effective expression method. When we used intron-mediated enhancement (IME) to improve circRNA expression of mouse Rtn4 (Nogo, a key protein in Nogo-Rho pathways) circRNA as a test case, we achieved a 4-6-fold improvement compared to an existing method. We further developed this approach into a general circRNA expression vector pCircRNA-DMo. An unexpected feature of our approach is its ability to promote translation of circRNA into detectable amounts of proteins. Intriguingly, both monomer and multimer repeating peptides can be observed as a result of rolling circle translation of RTN4 circRNA. We also confirmed the presence of both peptide forms in human and mouse brains, highlighting the significance of circRNA translation *in vivo*. In summary, we demonstrate the significant advantage of IME in enhancing circRNA biogenesis and hence our vector offers a robust platform for exploring potential circRNA peptide-encoding functions.

## Introduction

Circular RNAs (circRNA) are 5’ and 3’ covalently jointed isoforms from pre-mRNA back-splicing (1-3). They may play important roles in various biological processes (4-12). For example, Cdr1as/ciRS-7 (circular RNA sponge for miR-7) can bind miR-7 and regulate its activity (13-15). The function of circRNAs in cancer is also increasingly recognized (16-18).

Over-expression of circRNA is achieved by constructing inverted repeats in the 5’ and 3’ flanking intron sequences of the circularized exon (13,19,20). Since circularization competes with classic splicing, over-expression of circRNA is typically less efficient than for linear forms (21-23). Moreover, the accumulation of large amounts of linear precursor without efficient splicing may disturb circRNA-specific expression (24).

Intron-mediated enhancement (IME) is a conserved phenomenon across eukaryotes, including plants and mammals, where the expression of a gene is enhanced by an intron, often located upstream, and close to the start of transcription (25-31). Although the precise mechanism of IME is still largely unknown, the enhancement of mRNA accumulation from an intron-containing gene is often highly significant and is frequently used as a powerful method to increase gene expression, especially in transgenic organisms (30,32). However, it is unknown whether IME can also enhance circRNA expression.

Reticulon-4, also known as neurite outgrowth inhibitor (Nogo-A, B, C), encoded by the RTN4 gene, is an inhibitor of neurite outgrowth in the central nervous system of higher vertebrates, and may play critical role in blocking neuroregeneration after brain injury (33,34). RTN4 circRNA comprising exon 2 and 3 of the RTN4 gene is expressed in both human and mouse brain at detectable levels (4), thus representing a good model for studying circRNA expression.

We generated an IME circRNA expression vector and used it to increase the expression levels of Rtn4 circRNA in cell lines. Furthermore, we confirmed that such a vector is a general method for enhancing circRNA biogenesis by testing two additional IME introns. Finally, we show that boosted circRNA can be translated into proteins, demonstrating the significance of this method for studying circRNA function.

## Material and methods

### Plasmid construction

To construct the Rtn4 circRNA expression plasmid, Rtn4 circRNA exon genomic region (chr11: 29, 704, 497-29, 708, 881, mouse GRCm38/mm10) along with the partial 5’ and 3’ flanking intronic sequences (1014 nt and 111 nt) were amplified from N2a cell genomic DNA and inserted under the CMV promoter in pCMV-MIR (OriGene) vector. The resulted construct is named as control-2 (Fig. 1A). 800 nucleotides of 5’ intronic regions (chr11:29,704,521-29,705,320) of control-2 were reverse complementary inverted and added to 3’ flanking intronic region to promote the back-splicing (4,13). As the flanking introns lacks the 5’ and 3’ splice sites respectively, they would not be able to perform classic splicing to generate liner mRNA. The resulting construct was called pCircRNA-BE-Rtn4. To generate completely identical circRNA without additional sequences from restriction endonuclease sites in the circRNA expression vector, we performed overlap PCR instead of using restriction endonucleases in the plasmid construction. pCircRNA-DMo-Rtn4 was constructed based on pCircRNA-BE-Rtn4 with the addition of a chimeric intron from the pCI-neo-FLAG vector to the upstream of circRNA expression region under the same CMV promoter.

To create general circRNA expression vectors, multiple restriction endonuclease sites (BglII, NheI, BmtI, EcoRV, NotI, SacII, XbaI) were added to the original Rtn4 circRNA exon of pCircRNA-BE-Rtn4 or pCircRNA-DMo-Rtn4, producing the vectors for other circRNA expression, called pCircRNA-BE or pCircRNA-DMo (Supplementary Fig. 1).

The chimeric intron of pCircRNA-DMo-Rtn4 was replaced with the IVS1 intron or PAT1 intron 1 to generate pCircRNA-IVS1-Rtn4 and pCircRNA-PAT1-Rtn4 constructs (25,31). The linear mRNA expression of Rtn4 exon 2-exon3 was achieved by insertion of mouse Rtn4 exon 2-intron 2-exon 3 into pCMV-MIR vector under the CMV promoter and the resulting construct was called pCMV-Rtn4-Exon2-Exon3. A FLAG tag (DYKDDDDKPP) and stop codon (TGA) were added to the middle of exon 2 of pCircRNA-DMo-Rtn4 plasmid to form pCircRNA-DMo-Rtn4-FLAG and pCircRNA-DMo-Rtn4-Stop.

For pCircRNA-DMo-Rtn4-FLAG-ac was made by addition of ac dinucleotides after FLAG tag of pCircRNA-DMo-Rtn4-FLAG vector. To avoid too early termination in the second-round translation after the ac addition, the 7^th^ nucleotide (T) after the start codon (ATG) was changed to C.

Details of the oligos used in plasmid constructions are provided (Supplementary table). Plasmid DNAs were purified with an EndoFree Plasmid Maxi Kit (QIAGEN). To avoid inverted repeats mediated DNA recommendation in the circRNA expression vectors, all constructs were verified by double restriction endonuclease digestion and DNA Sanger sequencing (data not shown).

## Cell culture and plasmid DNA transfection

N2a, HeLa, HEK293 were cultured in Dulbecco’s modified Eagle medium (DMEM, Invitrogen), supplemented with 10% foetal bovine serum (Gibco), 10 mM sodium pyruvate (Sigma), 100 U/ml penicillin and 100 U/ml streptomycin (Gibco) at 37 °C in 5% (v/v) CO_2_. N2a-swe.10 cells were cultured in special medium described previously (35).

For transfection, 2.5 μg of plasmid DNA diluted in 150 μl Opti-MEM (Invitrogen) was mixed with 5 μl lipofectamine 2000 diluted in 150 μl Opti-MEM and the resulting transfection mix was added to about 0.5 million cells in 6 well plates. After 24 hours, the transfection mix was transferred to fresh DMEM medium. After 3 to 6 days of transfection, cells were collected for total RNA isolation and total protein extraction.

GFP fluorescence expressed from SV40 promoter in pCMV-MIR backbone was used as a reporter for transfection monitoring; for each circRNA expression plasmid transfection, almost equal fluorescence was observed. Moreover, GFP expression was measured by western blot with antibody against GFP (#2555, 1:1000, Cell signalling Technology). About equal GFP expressions were observed, demonstrating the almost same transfection efficiency for each circRNA expression plasmid DNA (Supplementary Fig. 2).

## Total RNA isolation and qRT-PCR

Total RNAs from N2a, N2a-swe.10, HeLa and HEK 293 cells were isolated using TRIzol reagent (Ambion) according to the manufacturer’s recommendations. 0.5 µg of total RNA was used to synthesize cDNA by random oligos with the SuperScript®III First-Strand Synthesis System (Invitrogen). Quantitative PCR amplification was performed using a 7900HT Fast Real Time PCR System (Applied Biosystems) using the Power SYBR Green PCR Master Mix (Applied Biosystems). Fold expression differences between treated samples versus control samples were calculated using the 2^−ΔΔ*C*^_T_ method with β-Actin mRNA as internal control (36).

### Northern blot

Northern blot was performed similarly as previously described (37). In brief, 15 µg of total RNA of HEK293 cells with transfection was separated in 1% agarose gel and transferred to positively charged nylon membrane (Amersham Hybond-N^+^, GE healthcare). DNA probe Rtn4-NB-R1 was labelled with ^32^P-phosphate group at 5’ and hybridized with the blot membrane at 42 °C.

Rtn4-NB-R1:5’TCCTGAACTAAATCTGGCGTTAGACCTTCAGGCATGGTTGCCACTACTGCCTCAGTCACC 3’

For RNase R treatment, 15 µg of total RNAs were digested with 10 unites of RNase R (RNR07250, Epicentre) for 1 hour at 37 °C.

### Western blot assays

Total protein from HEK293, N2a cells and mouse frontal cortex samples was prepared in RIPA buffer (50mM Tris-HCl pH 8.0, 150 mM NaCl, 0.1% (w/v) SDS, 0.5% (w/v) Na-Deoxycholate, 1% (v/v) NP40, 1* Roche cOmplete Protease Inhibitor and 1* PhosSTOP Phosphatase Inhibitor). Human brain samples were purchased from BioCat GmbH. 40 μg of total protein was fractioned on SDS-PAGE with reducing loading buffer (5% β-mercaptoethanol) and transferred to nitrocellulose membranes (Amersham), then immunoblotted against Nogo-A (Rtn4, 1:1000, #13401, Cell signalling Technology) and β-Actin (1: 15000; A5441, Sigma). Blots were then incubated with horseradish peroxidase conjugated secondary antibodies (goat-anti-rabbit, IgG (H+L), G21234; goat-anti-mouse IgG (H+L), G21040; Life technologies) and developed with ECL solution (Amersham) and imaged with the ChemiDoc MP Imaging System (Bio-Rad).

### Label-free quantitative proteomics

HEK293 cells individually transfected with pCircRNA-BE-Rtn4, pCircRNA-DMo-Rtn4 and the control empty plasmid were lysed and digested in solution with trypsin according to a previously established method (38). Briefly, cell pellets were heated and sonicated in lysis buffer (100 mM Tris-HCl, 6 M guanidinium chloride (guanidine hydrochloride, GuHCl), 10 mM TCEP (Tris (2-carboxyethyl) phosphine), 40 mM CAA (chloroacetamide)). After centrifugation, the diluted supernatant proteins were digested by trypsin (Promega, V5280) overnight and the resulting peptides were purified with C18-SD StageTip (38,39). The prepared peptides were analysed by an Orbitrap Fusion mass spectrometer (Thermo Fisher Scientific) with a nano-electrospray ion source, coupled with an EASY-nLC 1000 (Thermo Fisher Scientific) UHPLC. MaxQuant version 1.5.3.8 with an integrated Andromeda search engine was used to analyse the LC-MS/MS raw data (39,40). Detailed method is provided (Supplementary method).

### Immunoprecipitation and mass spectrometry

5 million of N2a with pCircRNA-DMo-Rtn4-FLAG transfection were collected, lysed and bound to the ANTI-FLAG-M2 affinity gel (FLAG Immunoprecipitation Kit, Sigma-Aldrich, FLAGIPT1) according to the technical bulletin. After washing, elution buffer of 5 ng/µl trypsin, 50 mM Tris-HCL, TCEP (tris(2-carboxyethyl) phosphine), 5 mM Chloroacetamide, pH 7.5 was added to the resin and incubated for 30 minutes at room temperature with gently shaking. The supernatant was transferred to a new tube and incubated at 37 ^º^C overnight to ensure a complete tryptic digestion. The rest procedure was performed as prescribed previously.

## Results

### Expression of Rtn4 circRNA using an existing method

A mouse Rtn4 circRNA expression cassette was amplified from N2a genomic DNA and constructed as previously described by Hansen et al. (13). It included 800-nucleotide reverse complementary repeats in the 5’ and 3’ flanking intronic regions and was inserted into the pCMV-MIR vector under the CMV promoter (Fig. 1A). Transfection of the resulting Rtn4 circRNA expression construct, pCircRNA-BE-Rtn4 (basal expression), into mouse neuroblastoma cell line (N2a) and its derivative, N2a-swe.10, followed by qRT-PCR with Rtn4 circRNA-specific PCR oligos (Rtn4-c-F and Rtn4-c-R, Fig. 1A) showed that Rtn4 circRNA over-expression was 3.9/5.8-fold higher than endogenous expression, i.e. after empty vector transfection (Fig. 1B). In two kinds of human cell lines, HeLa and HEK293, pCircRNA-BE-Rtn4 also expressed Rtn4 circRNA.

Moreover, to investigate whether any linear mRNA form is expressed from pCircRNA-BE-Rtn4, we performed northern blot with a DNA probe can detect both linear and circular forms of Rtn4 RNA. To indicate linear RNA migration on the gel, we transfected pCMV-Rtn4-Exon2-Exon3 which would express linear Rtn4-exon2-exon3 mRNA (Fig. 1A). As shown in and Fig. 1D, Rtn4 circRNA migrated faster than its linear counterpart in native agarose gel. Importantly, no linear Rtn4-exon2-exon3 mRNA was found in pCircRNA-BE-Rtn4 transfection (Fig. 1D), which was consistent with the design, as there was no intact intron at both 5’and 3’ for classic splicing. As negative control, construct lacking inverted repeat at 3’ flanking intronic region did not express any circRNA but produce linear RNA precursor (control-2 in Fig. 1A, D). Of note, as this construct (control-2) does not have intact 5’ flanking intron for both classic splicing and back-splicing, its precursor RNA (containing part of intron at 5’ side) may be processed by mRNA degradation pathway and would not be stable, thus it displayed a very weak band compared to the line of pCMV-Rtn4-Exon2-Exon3 in northern blot analysis (control-2 in Fig. 1A, D) (41,42). Furthermore, we used RNase R digestion to demonstrate the circularity of the expressed Rtn4 circRNA. As shown in Fig. 1D, the indicated band for Rtn4 circRNA still existed under RNase R treatment. As control, the linear Rtn4-exon2-exon3 mRNA disappeared under RNase R digestion (Fig. 1D).

Based on these results (especially the gold standard detection of circRNA by northern blot analysis with combination of RNase R treatment (37)) and the previous studies of circRNA biogenesis, we concluded that the flanking intronic sequences with inverted repeats can moderately promote Rtn4 circRNA biogenesis, thus serving as a good starting cassette for further optimizing circRNA vector design.

**Fig. 1.**
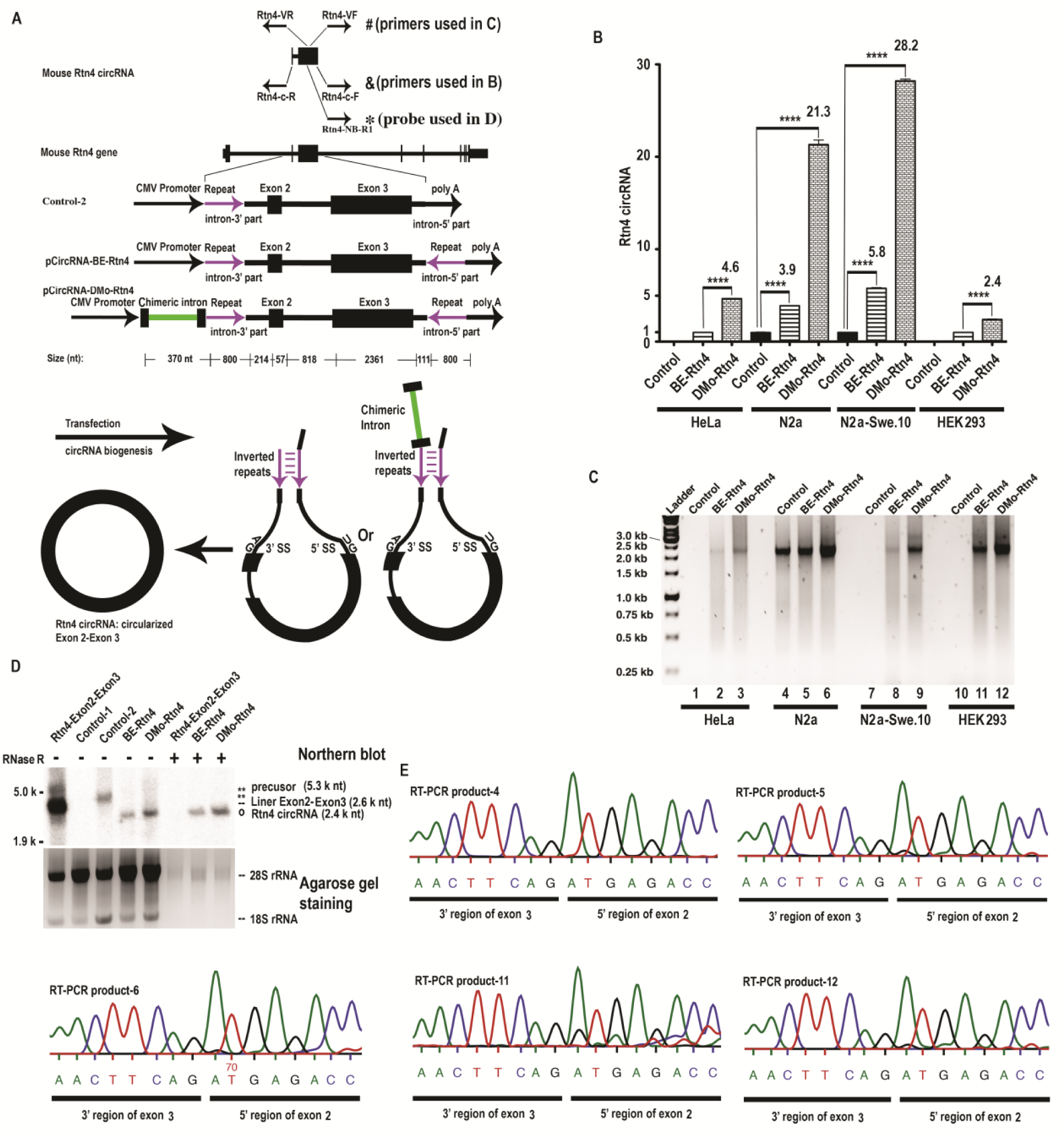
Mouse Rtn4 circRNA structure and expression in mammalian cells. A.Scheme of Rtn4 circRNA sequence localization in the Rtn4 gene and CircRNA expression cassette from pCircRNA-BE and pCircRNA-DMo vectors; mouse Rtn4 circRNA consists exon 2 and exon 3 of Rtn4 gene. 800 nts inverted repeats (purple colour) in the flanking introns were constructed to promote back-splicing through forming inter-intronic base-pairing; the flanking introns lacks 5’ and 3’ splice site respectively which would abolish the classic splicing of exon 2 and exon 3; chimeric intron was shown in green; & and #, the Rtn4 circRNA RT-PCR oligo positions; &, Rtn4-c-R, Rtn4-c-F were used in qRT-PCR to determine Rtn4 circRNA levels (results shown in B); #, Rtn4-VR, Rtn4-VF were used to verify Rtn4 circRNA back-splicing fidelity (results shown in C); *, the position of northern blot probe oligo (Rtn4-NB-R1) used in D. B.Rtn4 circRNA expression levels in transfected cells (HeLa, N2a, N2a-swe.10, HEK293 cell); Control = pCMV-MIR empty vector; BE-Rtn4 = pCircRNA-BE-Rtn4; DMo-Rtn4 = pCircRNA-DMo-Rtn4; All statistic T tests were performed by comparison with the control sample, *\*\*\**\*, P ≤ 0.0001, n ≥ 4; β-Actin mRNA was used as internal control. C.Agarose gel electrophoresis of RT-PCR products of Rtn4 circRNA to verify back-splicing fidelity (PCR primers: Rtn4-VR, Rtn4-VF); Control = pCMV-MIR empty vector; BE-Rtn4 = pCircRNA-BE-Rtn4; DMo-Rtn4 = pCircRNA-DMo-Rtn4. The entire un-spliced primary precursor transcript is about 5.3 k nt; the spliced circular form of Exon 2 and exon 3 without intron is 2.4 k nt. The size of Intron between two exons is 818 nt. The expected amplicon size is 2.4 kb. As the products migrated between 2.0 and 2.5 kb, proving that the internal intron was spliced out in Rtn4 circRNA biogenesis. lane 1-3, HeLa cells; lane 4-6, N2a cells; lane 7-9, N2a-swe.10 cells; lane 10-12, HEK293 cells. PCR products were sequenced and aligned (data not shown). D.Northern blot of Rtn4 circRNA in transfected HEK293 cells. Control-1 = pCMV-MIR empty vector; Rtn4-Exon2-Exon3 = pCMV-Rtn4-Exon2-Exon3; Control-2, the construct without inverted repeat at 3’ flanking intronic region; BE-Rtn4 = pCircRNA-BE-Rtn4; DMo-Rtn4 = pCircRNA-DMo-Rtn4; -, no RNase R treatment; +, with RNase R treatment; ethidium bromide staining of agarose gel showing 28S and 18S rRNAs were used as loading control. E.Sequencing of junction site of back-splicing of Rtn4 circRNA overexpressed in N2a and HEK293 cells. The RT-PCR products from C were sequenced and the junction regions were shown; RT-PCR product-4, 5, 6 were from N2a cell; RT-PCR product-11, 12 were from HEK293 cell.

### A neighbouring chimeric intron significantly enhances Rtn4 circRNA biogenesis: Intron-mediated enhancement (IME) in circRNA expression

The presence of an intron frequently increases mRNA accumulation in gene expression, described as intron-mediated enhancement (IME) (30,32). An interesting question is whether the neighbouring intron can also promote circRNA expression. To test this, we amplified a chimeric intron that has shown a strong IME effect in mRNA expression from pCI-neo vector (Promega) and inserted it upstream of the Rtn4 circRNA cassette under the same CMV promoter (Fig. 1A). The resulting vector was called pCircRNA-DMo-Rtn4.

Expression studies in N2a and N2a-swe.10 showed that Rtn4 circRNA biogenesis was significantly improved from 4-6-fold up to 21-28-fold, representing robust enhancement of circRNA expression (Fig. 1B). The IME effect on Rtn4 circRNA expression in the two kinds of human cell lines, HeLa and HEK293 resulted a 4.6/2.4-fold increase in Rtn4 circRNA expression in these cells with the chimeric intron vector (pCircRNA-DMo-Rtn4), compared to the intron-less control (pCircRNA-BE-Rtn4), thus demonstrating that the chimeric intron-containing circRNA expression vector drives ubiquitous enhancement effects in various cell lines (Fig. 1B). Our data clearly show that intron-mediated enhancement (IME) boosts circRNA expression, thus providing a good strategy for circRNA expression cassette construction. Of-note, as shown in the northern blot analysis of Rtn4 circRNA, pCircRNA-DMo-Rtn4 did not produce linear Rtn4-exon2-exon3 mRNA, which is consistent with its design (Fig. 1D). Furthermore, its expressed Rtn4 circRNA was resistance to RNase R treatment (Fig. 1D).

### Back-splicing fidelity of pCircRNA-BE-Rtn4 and pCircRNA-DMo-Rtn4

Next, we evaluated the back-splicing fidelity of our circRNA expression vectors. As shown above, the unique RT-PCR oligos (Rtn4-VF, Rtn4-VR in Fig. 1A) could amply the full sequence of Rtn4 circRNA. Agarose gel electrophoresis of the RT-PCR product showed that expressed Rtn4 circRNAs were all the same size (Fig. 1C). Further sequencing of the RT-PCR products confirmed that the Rtn4 circRNA sequences were identical to the wild type circRNAs without any sequence differences in either human or mouse cell lines (data not shown). Importantly, sequencing results also confirmed the junction sequences are presented in both wildtype and vector expressed Rtn4 circRNA in N2a and HEK293 cells (Fig. 1E).

We conclude that both of our expression vectors produce circRNA with the same sequence as the endogenous copy, representing high back-splicing fidelity.

### IVS1 and PAT1 introns also boost circRNA biogenesis: Ubiquitously robust enhancement by various introns

To further test whether the IME effect on circRNA expression is universal to different introns, we replaced the chimeric intron in pCircRNA-DMo-Rtn4 with two other introns, IVS1 and PAT1 intron 1, which were shown to stably enhance their respective mRNA expression (25,31). The resulting Rtn4 circRNA expression vectors, pCircRNA-IVS1-Rtn4 and pCircRNA-PAT1-Rtn4 (Fig. 2A) were transfected into N2a cells, and the qRT-PCR of Rtn4 circRNA showed that IVS1 and PAT1 introns enhance Rtn4 circRNA expression from 3.9-fold (pCircRNA-BE-Rtn4) to 7.8 and 6.8-fold, representing about 2-fold enhancement, although not as robustly as the ca. 20-fold enhancement driven by pCircRNA-DMo-Rtn4 (Fig. 2A, B).

Based on these results, we conclude that various introns could boost circRNA, demonstrating the universal existence of the IME effect in circRNA biogenesis.

**Fig. 2.**
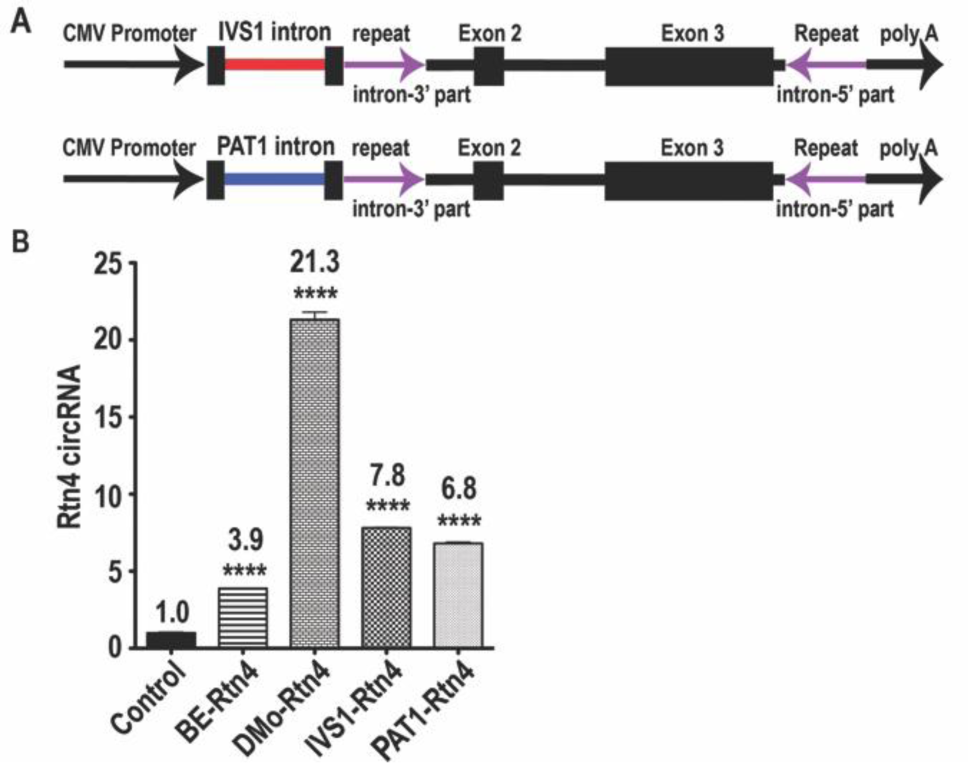
Expression of Rtn4 circRNA with IVS1 and PAT1 introns. A. Scheme of the Rtn4 circRNA expression cassette for pCircRNA-IVS1-Rtn4 and pCircRNA-PAT1-Rtn4 constructs. IVS1 intron was shown in red colour and PAT1 intron was shown in blue colour. B. Rtn4 circRNA expression in N2a cells after transfection with Rtn4 circRNA constructs. All statistic T tests were performed by comparison with the control sample, *\*\*\**\*, P ≤ 0.0001,*\*\**\*, P ≤ 0.001, n ≥ 4.

### Protein product from Rtn4 circRNA

To investigate the coding potential of circRNAs, we performed Western blots using Anti-Nogo-A antibody to detect proteins in the Rtn4 circRNA overexpressing cells. Since there was very weak / no expression of endogenous RTN4 protein in HEK 293 cells, we chose these cell extracts as controls for the Rtn4 circRNA translation study. As a positive control, we used pCMV-Rtn4 exon 2-3 (Fig. 3A), which expresses a linear counterpart mRNA of the Rtn4 circRNA exons. The translated protein from Rtn4 exon 2-3 mRNA migrated at around 150 kDa in the Western blot (Fig. 3C). The pCircRNA-BE-Rtn4 product failed to produce detectable protein, but pCircRNA-DMo-Rtn4 not only clearly expressed a Rtn4 related protein of around 150 kDa in HEK293 cells, but interestingly also a much larger form, possibly due to repeating peptides expressed from continuous translation, as discussed below (Fig. 3B). These results mirror the dramatic induction of expression of Rtn4 circRNA due to the IME effect.

**Fig. 3.**
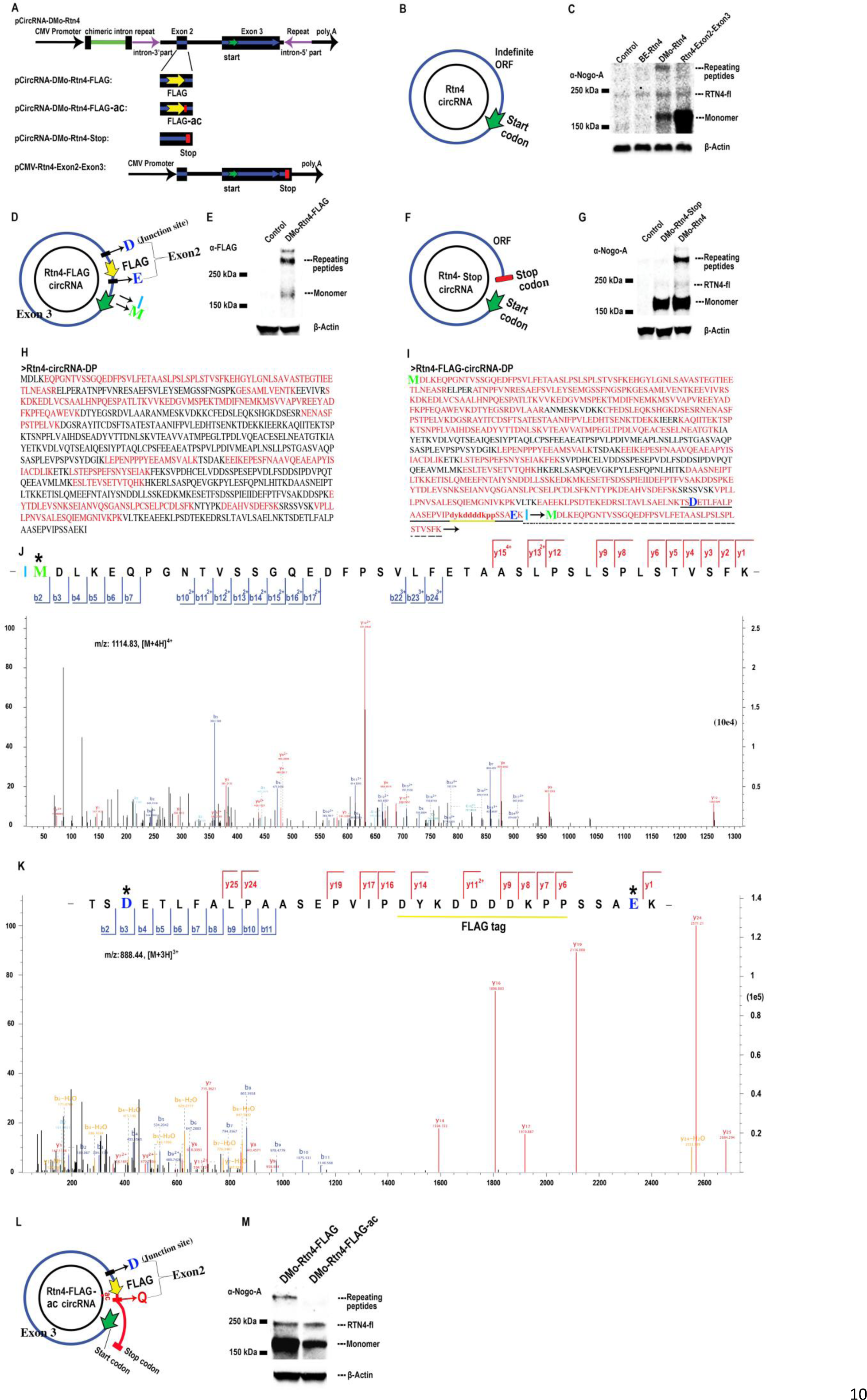
Rolling circle translation of Rtn4 circRNA. A.Scheme illustrating the insertion of a FLAG tag and stop codon into exon 2 of the open reading frame of Rtn4 circRNA. Blue, open reading frame; green arrow, start codon, the first AUG in the ORF; yellow arrow, FLAG tag in the pCircRNAsDMo-Rtn4-FLAG construct; red rectangle, stop codon (UGA) in the pCircRNA-DMo-Rtn4-Stop construct. B. Indefinite open reading frame (ORF) of Rtn4 circRNA. Start codon, AUG in the ORF. C. Western blot of Rtn4 circRNA translated proteins in HEK293 cell detected by antibodies against Nogo-A (α-Nogo-A). Control, empty vector; BE-Rtn4 = pCircRNA-BE-Rtn4; DMo-Rtn4 = pCircRNA-DMo-Rtn4; Rtn4-Exon2-Exon3 = pCMV-Rtn4-Exon2-Exon3. On the right repeating peptides are multimers of the Rtn4 circRNA translation product; RTN4-fl indicates the endogenous RTN4 full length protein (Nogo-A); monomer was the single round translation product of Rtn4 circRNA. The calculated MW of monomer is 88.2 kDa. As it has very acidic pI (4.3), which may cause it migrate larger to 150 kDa position (43). D. Open reading frame of Rtn4-FLAG circRNA. The aspartic acid (blue D) is translated from the junction site of back-splicing; FLAG tag was inserted in exon 2; the blue E is the translated from junction site of exon2-exon3; the green M is translated from start codon. E. Western blot of pCircRNA-DMo-Rtn4-FLAG transfected N2a cells using an anti-FLAG antibody (α-FLAG). Control, the empty vector; DMo-Rtn4-FLAG = pCircRNA-DMo-Rtn4-FLAG; off-note, compared to C, the intensity of repeating peptides is higher than monomer; the reason is unknown, possibly due to the different antibody affinity between multimer and monomer. F. Open reading frame of Rtn4-stop circRNA; red rectangle, stop codon (UGA). G. Western blot of pCircRNA-DMo-Rtn4-Stop transfected HEK293 cells using an anti-Nogo A antibody (α-Nogo-A). Control, the empty vector; DMo-Rtn4-Stop = pCircRNA-DMo-Rtn4-Stop; DMo-Rtn4 = pCircRNA-DMo-Rtn4. H. Mass spectrometry results of Rtn4 circRNA derived peptides (Rtn4-circRNA-DP) expressed in HEK293 cell. The predicted sequence of monomer mouse Rtn4 circRNA derived peptides; red sequences, the detected peptides (detailed peptide spectrometry was provided in Supplementary Fig. 3). I. Mass spectrometry results of immunoprecipitation of Rtn4-FLAG circRNA derived peptides (Rtn4-FLAG-circRNA-DP) expressed in N2a cell. Black sequences, undetected peptides; 76.8% amino acids are detected in mass spectrometry (shown in red); Blue D was the translated amino acid from the junction site of back-splicing, which is the direct evidence of translation from Rtn4-FLAG-circRNA; blue E is the translation product of junction site formed after the removal of the internal intron between exon2 and exon3; light blue I is the last amino acid of one round transition; green M is from the start codon (AUG); peptides labelled with underline were the junction peptides; detected peptides with dotted line was the direct evidence of rolling circle translation of Rtn4-FLAG-circRNA; arrow indicates continuous repeating translation. J. Mass spectrometry of peptide labelled with underline in I; the direct evidence of the protein translation from the junction site of back-splicing of Rtn4. K. Mass spectrometry of peptide labelled with dotted line in I; the direct evidence of the continuous translation of Rtn4 circRNA which generate repeating peptides. L. Open reading frame of Rtn4-FLAG-ac circRNA; ac, adenosine and cytosine; red rectangle, stop codon (UAA). M. Western blot analysis of pCircRNA-DMo-Rtn4-ac transfected in N2a cells using an anti-Nogo A antibody (α-Nogo-A). DMo-Rtn4-FLAG = pCircRNA-DMo-Rtn4-FLAG; DMo-Rtn4-FLAG-ac = pCircRNA-DMo-Rtn4-FLAG-ac.

### The indefinite translation of Rtn4 circRNA

Examination of the open reading frame of Rtn4 circRNA found that this circRNA does not have a stop codon, so represents a circular ORF that may produce repeating peptides, with potentially very high molecular weight. As predicted, we observed a high molecular weight band in the western blot against Rtn4 protein in pCircRNA-DMo-Rtn4 transfected HEK293 cells under reducing condition (Fig. 3C). These large proteins presumably arise from Rtn4 circRNA continuous translation (Fig. 3B). To further verify the specificity of these large proteins, and that they resulted from circRNA translation, we added a FLAG tag to the 5’ part of Rtn4 circRNA to generate pCircRNA-DMo-Rtn4-FLAG (Fig. 3D). Since the FLAG tag was located upstream of the start codon in the construct, FLAG can only be expressed from circular translation of Rtn4 circRNA (Fig. 3A, D). Western blotting for the FLAG tag detected both the monomer and high molecular weight bands of large proteins, further suggesting the continuous translation of the circular ORF of Rtn4 circRNA (Fig. 3A, D, E). As these large proteins with high molecular weight may also rise from protein aggregation of the monomer (although with less possibility), we then added a stop codon into the middle of exon 2 (pCircRNA-DMo-Rtn4-Stop, Fig. 3A, F), Western blot analysis with anti-Nogo-A antibody revealed that the stop codon abolished these high molecular bands (DMo-Rtn4-Stop in Fig. 3G), while the monomer of Rtn4 circRNA translated protein remained. If these high molecular weight bands were from protein aggregation, the stop code addition should not affect their migration. Thus, the abolishing of high molecular bands by stop codon addition in the ORF solidly confirmed the expression of repeating peptides from the continuous translation of Rtn4 circRNA.

To further investigate the identity of the peptides derived from Rtn4 circRNA translation, we performed shotgun mass spectrometry of Rtn4 peptides overexpressed in HEK293 cells. We detected 17 unique peptides (Supplementary Fig. 3), which covered about 38% of the putative translated monomer protein sequence from Rtn4 circRNA (the red sequences in Fig. 3H,). Since the putative protein sequence from mouse Rtn4 circRNA translation is significantly different from the counterpart human RTN4 protein fragment (Supplementary Fig. 4), we could confirm that peptides detected by mass spectrometry were derived from mouse Rtn4 circRNA translation.

To further prove the continuous translation of Rtn4 circRNA, we performed immunoprecipitation (IP) of Rtn4-FLAG circRNA derived proteins with FLAG antibody. Mass spectrometry (MS) of the immunoprecipitated proteins revealed the junction peptides (Fig. 3D, I, J, K). 76.8% amino acids were detected in this mass spectrometry (Fig. 3I), representing an enrichment of FLGA-tagged immunoprecipitation. The aspartic acid translated from the junction site of back splicing (the blue D in Fig. 3D) was observed in the received spectrometry (the blue D in Fig. 3K), thus providing the definitive evidence of Rnt4 circRNA translation.

Moreover, we also observed the junction region of repeating peptides. As shown in Fig. 3, the last amino acid residue (isoleucine, I) and the first amino acid residue (methionine, M) of one round translation was presented in the single peptide spectrometry (Fig.3D, the dotted line labelled peptides in Fig.3I, Fig. 3J), which can be only formed through continuous translation of Rtn4 circRNA, thus providing the direct evidence of rolling cycle translation.

Finally, we add two additional nucleotides (ac) after the FALG tag of pCircRNA-DMo-Rtn4-FLAG (Fig. 3A, L, M). The resulted plasmid was named as pCircRNA-DMo-Rtn4-FLAG-ac and its generated circRNA was called as Rtn4-FLAG-ac circRNA. The addition of ac dinucleotides to Rtn4-FLAG circRNA would cause frame shift of the open reading frame and it will generate 69 new amino acids at carboxyl terminal (Fig. 3A, L, M) until the stop codon (UAA), thus rolling cycle translation would be abolished. Indeed, western blot analysis of N2a cells with pCircRNA-DMo-Rtn4-FLAG-ac transfection clearly revealed only single monomer band existed and the multimer repeating peptide was dismissed (Fig. 3A, L, M).

In summary, we conclude that Rtn4 circRNA is translated into monomer and multimer repeating peptides. Similarly, we observed that the chimeric intron could induce Rtn4 circRNA production and hence elevated translation in N2a cells (Supplementary Fig. 5). Furthermore, IVS1 and PAT1 introns could also elevate both the expression level of Rtn4 circRNA and its translated protein in N2a cells (Supplementary Fig. 5). Taken together, we conclude that IME could significantly enhance gene expression of circRNAs.

### *In vivo* translation of endogenous RTN4 circRNA in human and mouse brain

Since RTN4 circRNA has been detected in human and mouse brain, we then asked whether it is translated *in vivo*. Western blot analysis of human and mouse brain tissues using an anti-Nogo-A antibody detected both the monomer and repeating peptides from RTN4 circRNA translation along with the full length RTN4 protein (Fig. 4), demonstrating the indefinite translation of RTN4 circRNA *in vivo*.

**Fig. 4.**
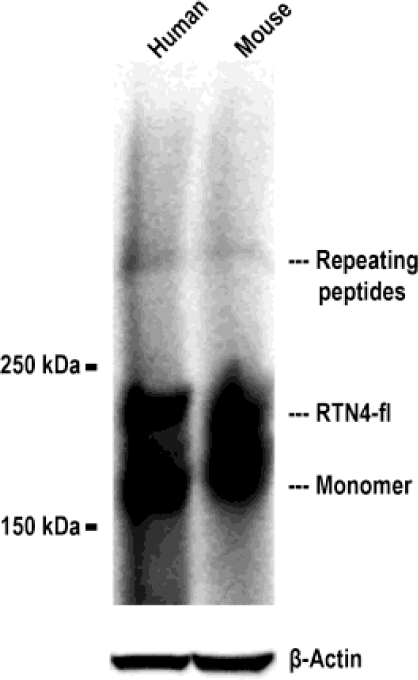
RTN4 circRNA-derived peptides expressed in human and mouse brain. Human brain frontal lobe and mouse frontal cortex samples were fractioned by SDS-PAGE and immunoblotted against Nogo-A (α-Nogo-A). RTN4-fl, RTN4 full length protein (Nogo-A); monomer, monomer of RTN4 circRNA derived peptide; repeating peptides, multimer of RTN4 circRNA derived peptide. β-Actin was used as a loading control.

## Discussion

Perhaps the most intriguing question about circRNAs concerns their functions. Although several hypotheses have been proposed, the biological roles of most circRNAs are still elusive. Previous studies on circRNA functions have been hampered by the lack of an efficient molecular tool to alter their expression levels. Using three different neighbouring introns, we clearly demonstrate that intron-mediated enhancement (IME) can boost circRNA expression. Interestingly, since the majority of highly expressed circRNAs are derived from the middle exon of host gene (19), their flanking introns may have IME effects that promote corresponding circRNA expression. Here, we propose that IME represents a dramatically improved method to accelerate studies into circRNA function.

CircRNAs are mostly hosted by protein encoding genes and produced by back-splicing in parallel to the linear counterpart mRNA splicing. Since these RNA species are located mostly in the cytoplasm, an attractive and immerging idea is that circRNAs may serve as templates for protein synthesis.

Previous studies have shown that several *in vitro* synthesized circular RNAs can be translated *in vitro* and in cells (44,45). An artificial split GFP circRNA can produce intact GFP protein (46). Furthermore, it has been shown that endogenous circ-ZNF609 circRNA can be translated into protein in a cell line (47). A circRNA generated from the muscleblind locus is reported to encode a protein in a drosophila fly’s head (48). N6-methyladenosine (m6A) modification of circRNA could induce efficient initiation of protein translation of circRNAs (49). These findings highlight the possibility that many, if not most circRNAs are translatable, hence acting similarly as their linear counterpart mRNA (50,51).

Here, we show that IME can significantly elevate Rtn4 circRNA production, thus offering a better expression tool for investigating the functions of circRNAs, especially protein synthesis. We have recently used pCircRNA-DMo vector to express two other circRNAs in cell lines. Preliminary results show that IME can significantly improve their expressions (Dingding Mo, unpublished data).

Interestingly, we demonstrate that Rtn4 circRNA is translated into monomer and multimer repeating peptide, representing rolling circle translation of an indefinite ORF. Importantly, the expression of both monomer and repeating peptides from RTN4 circRNA translation in human and mouse brain highlight their significance *in vivo*. Such unique rolling cycle translation had previously been described in an artificial circRNA (44,45). Here, we confirm its existence in natural circRNA. Such a finding suggests that other circRNAs may also have indefinite ORFs, thus adding new dimensions to RNA translation and protein diversity. (52). Interestingly, peptide species from the indefinite ORF of RTN4 circRNA are limited (Fig. 3), indicating that there is an unknown mechanism to terminate the translation from the circular RNA even without the stop codon. It will be interesting to address deeply about how translation of circRNA is initiated by ribosome and how it is terminated.

So far, we do not know whether such RTN4 circRNA-derived proteins are functionally important. Since RTN4 protein is an important neurite outgrowth inhibitor and a key protein in Nogo-Rho pathways, antagonizing RTN4 full-length protein is one of the approaches being investigated in neuroregenerative medicine (53). Since the second exon of the RTN4 circRNA encodes theΔ20 domain mediating neurite growth inhibition (33), RTN4 circRNA-derived proteins containing theΔ20 domain may also inhibit neurite growth and represent new players in Nogo-Rho pathways.

In summary, we conclude that IME provides robust benefits for circRNA gene expression, thus serving as an excellent tool to investigate various circRNA functions. However, further study is required to decipher the mechanism of how IME boosts circRNA expression and subsequent translation. Specifically, in terms of RTN4 circRNA, it may prove to be a clinically relevant tool to investigate the function of RTN4 circRNA-derived proteins.

## Acknowledgements

The authors acknowledge the department of biological mechanisms of ageing led by Linda Partridge for sharing chemicals and instruments. The authors also acknowledge the sharing of HeLa, HEK293 cells from Nils-Göran Larsson’s group. pCI-neo-FLAG is gift from Niels Gehring (University of Cologne). N2a-swe.10 cell line was kindly provided by Gopal Thinakaran (University of Chicago). The mouse sample used in this study was from the transgenic core facility of the host institute. The authors also appreciate Adam Antebi and Gabriella B Lundkvist for reading the manuscript.

## Author Contributions

DM designed and conceived the study. DM performed the experiments, analysed the results, prepared the figures, and wrote the manuscript. XL performed the mass spectrometry experiment. DC and JV participated in plasmid constructions.

## Supplementary method

### Label-free quantitative proteomics

HEK293 cells individually transfected with pCircRNA-BE-Rtn4, pCircRNA-DMo-Rtn4 and the control empty plasmid were lysed and in-solution digested with trypsin according to the previously established method (1). Briefly, Cell pellets were heated and sonicated in lysis buffer (100 mM Tris-HCl, 6 M guanidine hydrochloride, 10 mM TCEP (Tris (2-carboxyethyl) phosphine), 40 mM CAA (chloroacetamide)). After centrifugation, the diluted supernatant was diluted and digested with trypsin (Promega, V5280) overnight and the resulted peptides were purified with C18-SD StrageTip (1,2).

The peptides were analyzed using an Orbitrap QExactive HF mass spectrometer (Thermo Fisher Scientific) with a Nano-electrospray ion source, coupled with an EASY-nLC 1000 (Thermo Fisher Scientific) UHPLC. A 25-cm long reverse-phase C18 column with 75 μm inner diameter (PicoFrit, LC Packings) was used for separating peptides. The LC run lasted 150 min with a concentration of 6% solvent B (0.1% formic acid in 80% acetonitrile) increasing to 31% over 155 min and further to 50% over 20 min. The column was subsequently washed and re-equilibrated. MS spectra were acquired in a data-dependent manner with top 15. For MS, the mass range was set to 300-1800m/z and resolution to 60K at 200m/z. The AGC target of MS was set to 3e6, and the maximum injection time was 80 ms. Peptides were fragmented with HCD with collision energy of 27. The resolution was set to 15K. The AGC target of MSMS was 1e5 and the maximum injection time was 80 ms.

For mass spectrometric analysis of immune-precipitated peptides proteins, peptides were separated on a 25cm long, 75 mm internal diameter C18 column using an EASY-nLC 1200 (Thermo Fisher Scientific, Germany) as before. The gradient is from 0% to 6% buffer B for 2 min, from 6% to 25% buffer B for 60 min, from 25% to 50% buffer B for 20 min at 200 nl/min. Eluting peptides were analyzed on Orbitrap Fusion mass spectrometer (Thermo Fisher Scientific. Peptide precursor mass to charge ratio (m/z) measurements (MS1) were carried out at resolution of 60000 in the range of 300 to 1500 m/z. Cycle time of 3 sec was used for selecting the most intense precursors with charge state from 2 to 7 for HCD fragmentation with normalized collision energy of 25. The m/z of the peptide fragments (MS2) were measured at resolution of 30000 using an AGC target of 1e5 and maximum injection time100 ms. Upon fragmentation precursors were put on an exclusion list for 45 secs

MaxQuant version 1.5.3.8 with integrated Andromeda search engine was used for analyzing the LC/MSMS raw data (2,3). The raw data were searched against the mouse proteome from UniProt (knowledgebase 2016_04). Following parameters were used for data analysis: for ‘’fixed modification’’: cysteine carbamidomethylation, methionine oxidation; for ‘’variable modification’’: methionine oxidation and protein N-terminal acetylation; for ‘’digestion’’ specific with Trypsin/P, Max. missed cleavages 2; for label-free quantification, match between runs is selected. Other parameters were set as default.

Protein quantification significant analysis was performed with the Perseus statistical framework (http://www.perseus-framework.org/) version 1.5.2.4. After removing the contaminants and reverse identifications, the intensities were transformed to log2. Two-sample test was performed to identify the differentially expressed proteins. Proteins with a p-value of less than 0.05 were designated as significantly differentially expressed.

## Supplementary table

Oligos used in this study

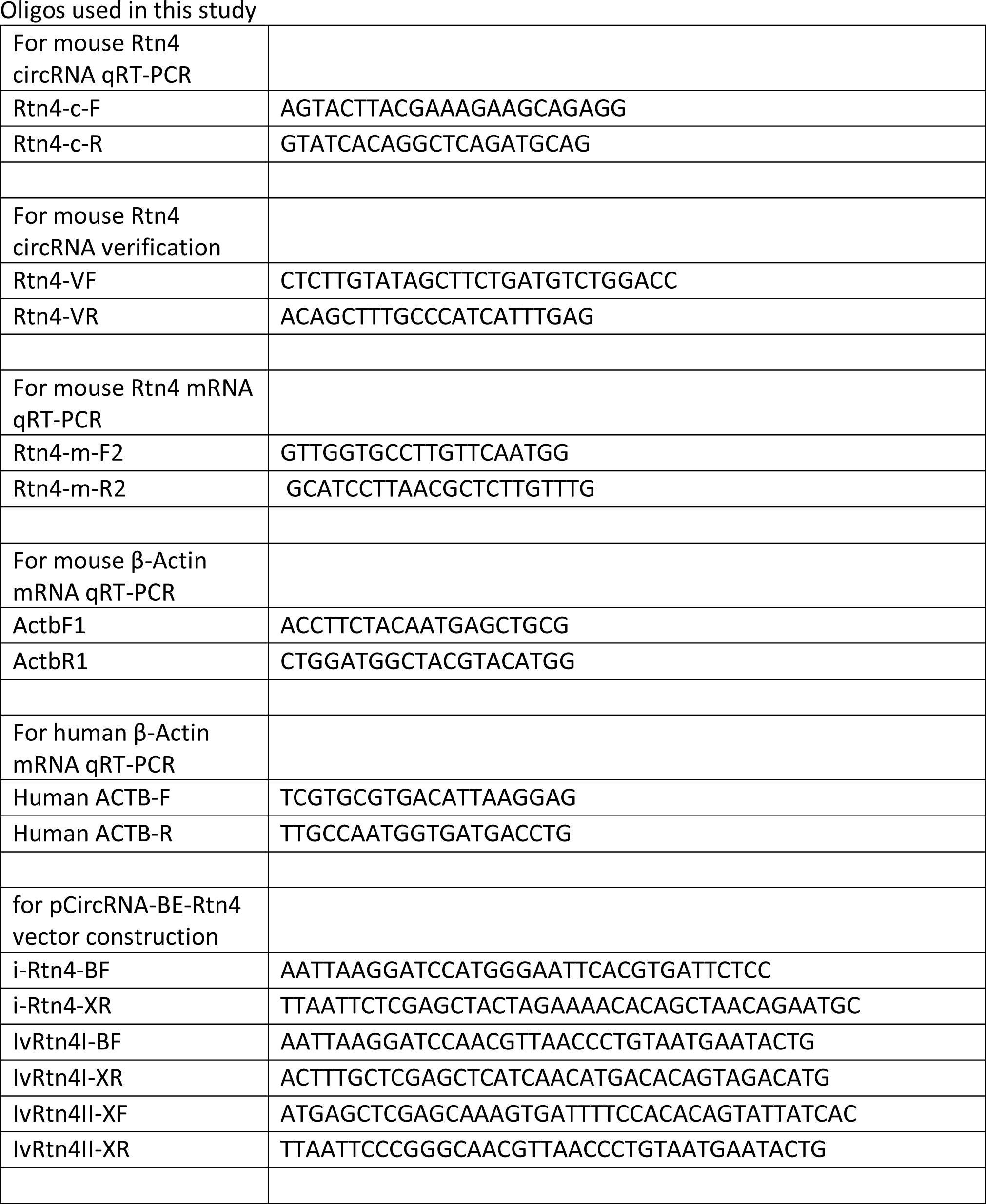
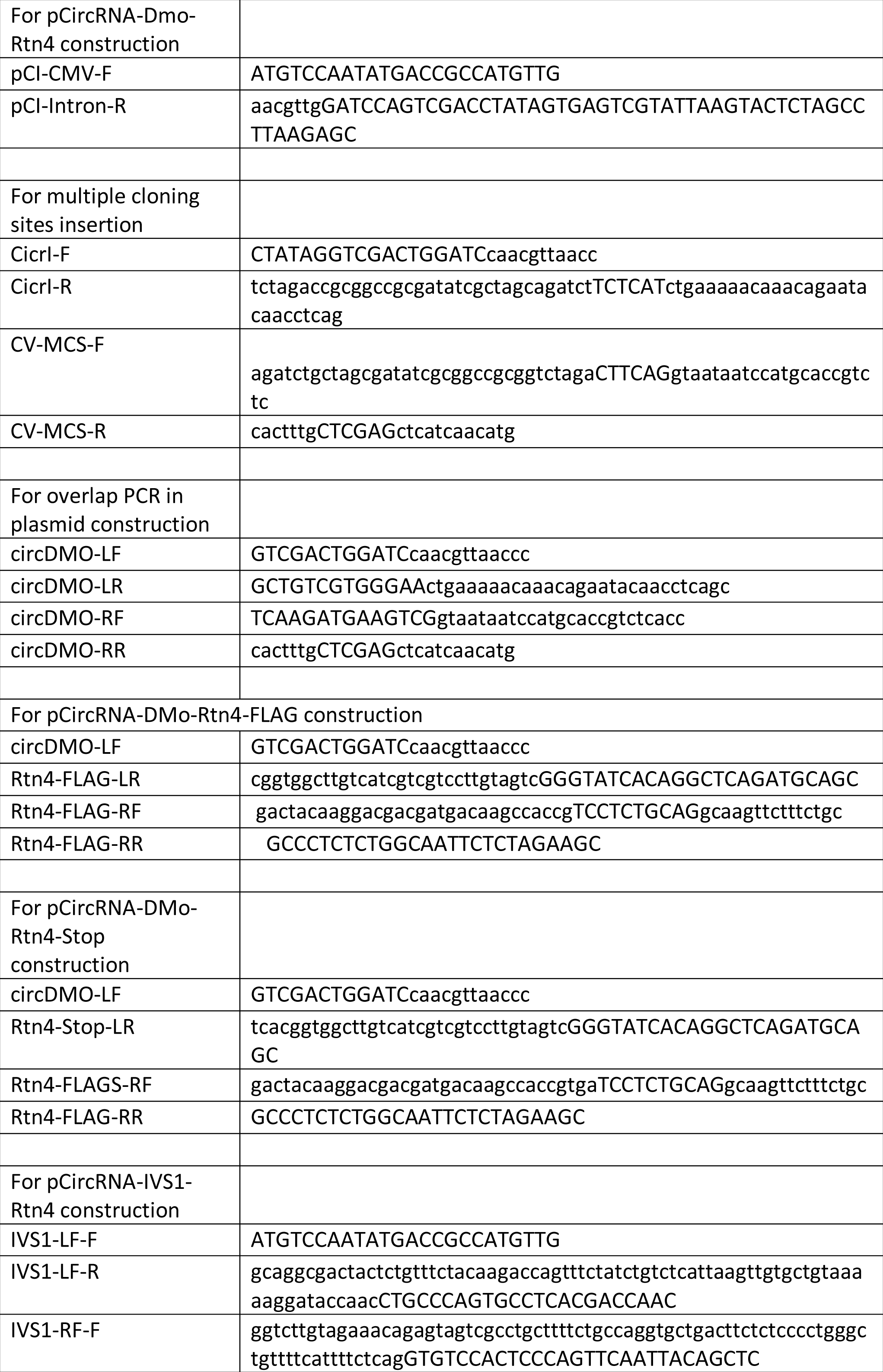
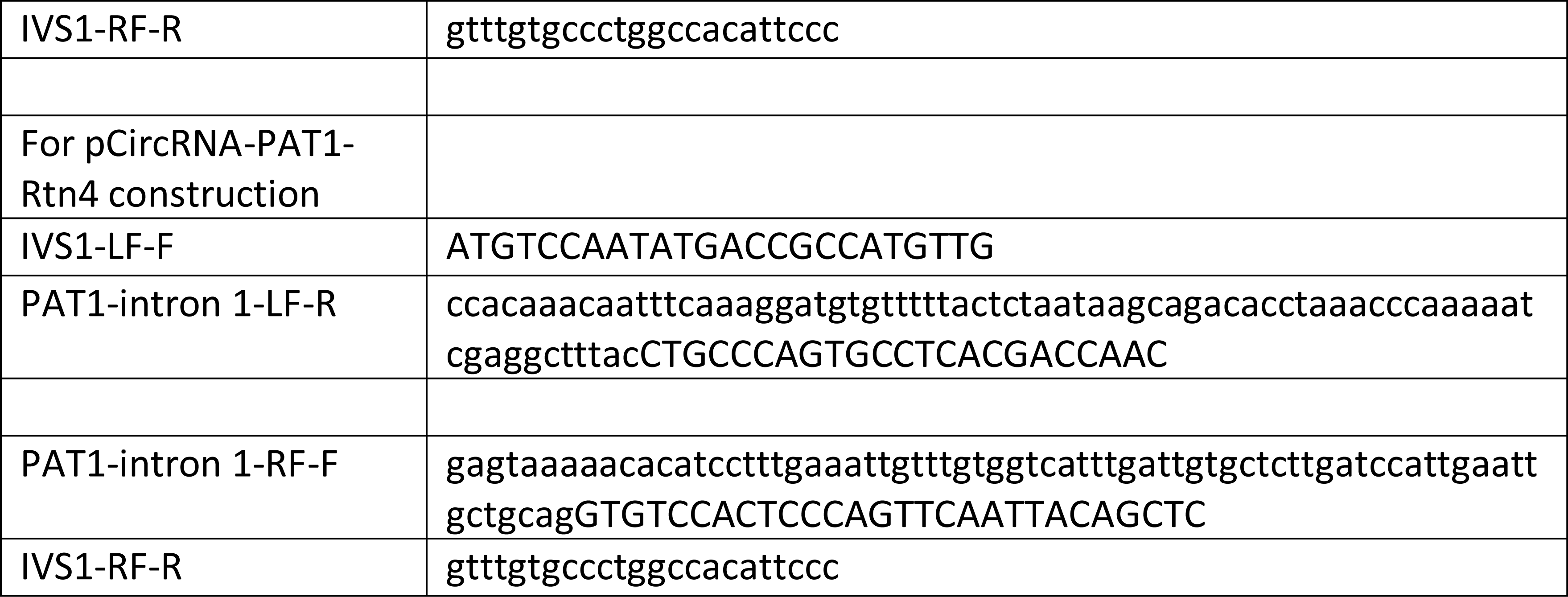

## Supplementary figures

**Supplementary Fig. 1.**
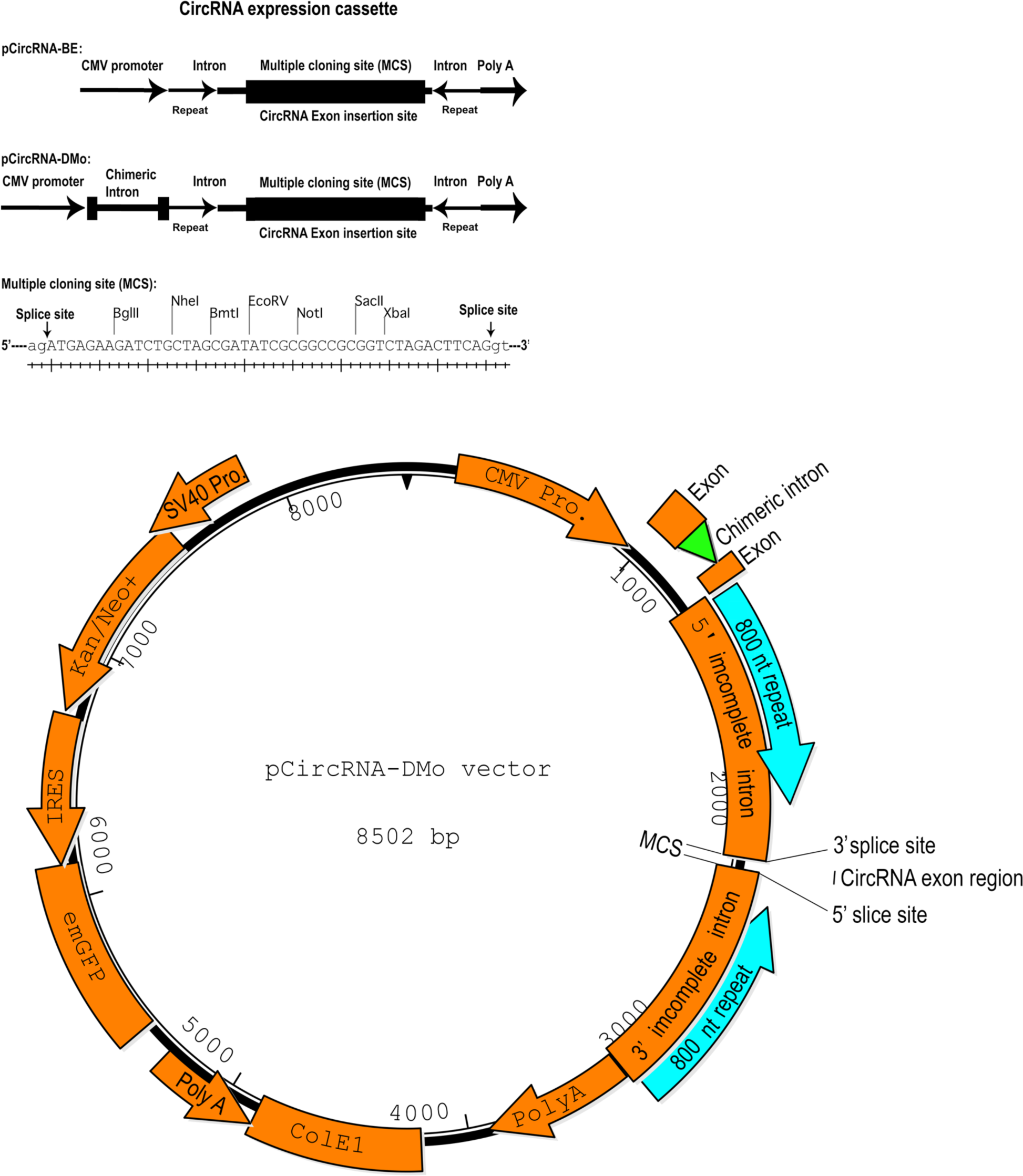
CircRNA expression cassette of pCircRNA-BE and pCircRNA-DMo vectors. Multiple cloning sites (MCS) contain following restriction endonuclease sites: BglII, NheI, BmtI, EcoRV, NotI, SacII, XbaI. 5’ and 3’ splice site is indicated by arrow. tmGFP in the original pCMV-MIR vector was changed to emGFP.

**Supplementary Fig. 2.**
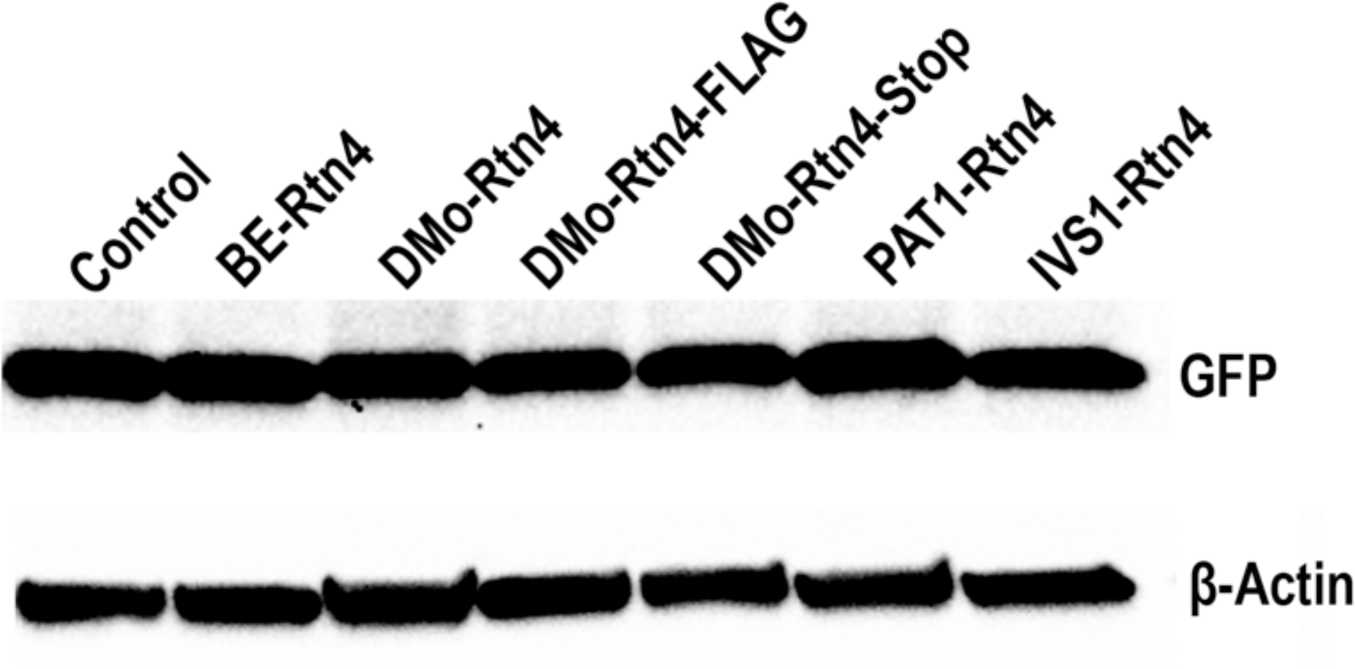
Equal transfection efficiency from CircRNA expression vectors. GPP expression from SV40 promoter (Supplementary Fig. 1) in each circRNA expression plasmid DNA transfection in HEK293 cell was measured by western blot analysis with antibody against GFP. β-Actin was used as loading control. Control, pCircRNA-DMo; BE-Rtn4, pCircRNA-BE-Rtn4; DMo-Rtn4, pCircRNA-DMo-Rtn4; DMo-Rtn4-FLAG, pCircRNA-DMo-Rtn4-FLAG; DMo-Rtn4-Stop, pCircRNA-DMo-Rtn4-Stop; PAT1-Rtn4, pCircRNA-PAT1-Rtn4; IVS1-Rtn4, pCircRNA-IVS1-Rtn4.

**Supplementary Fig. 3.**
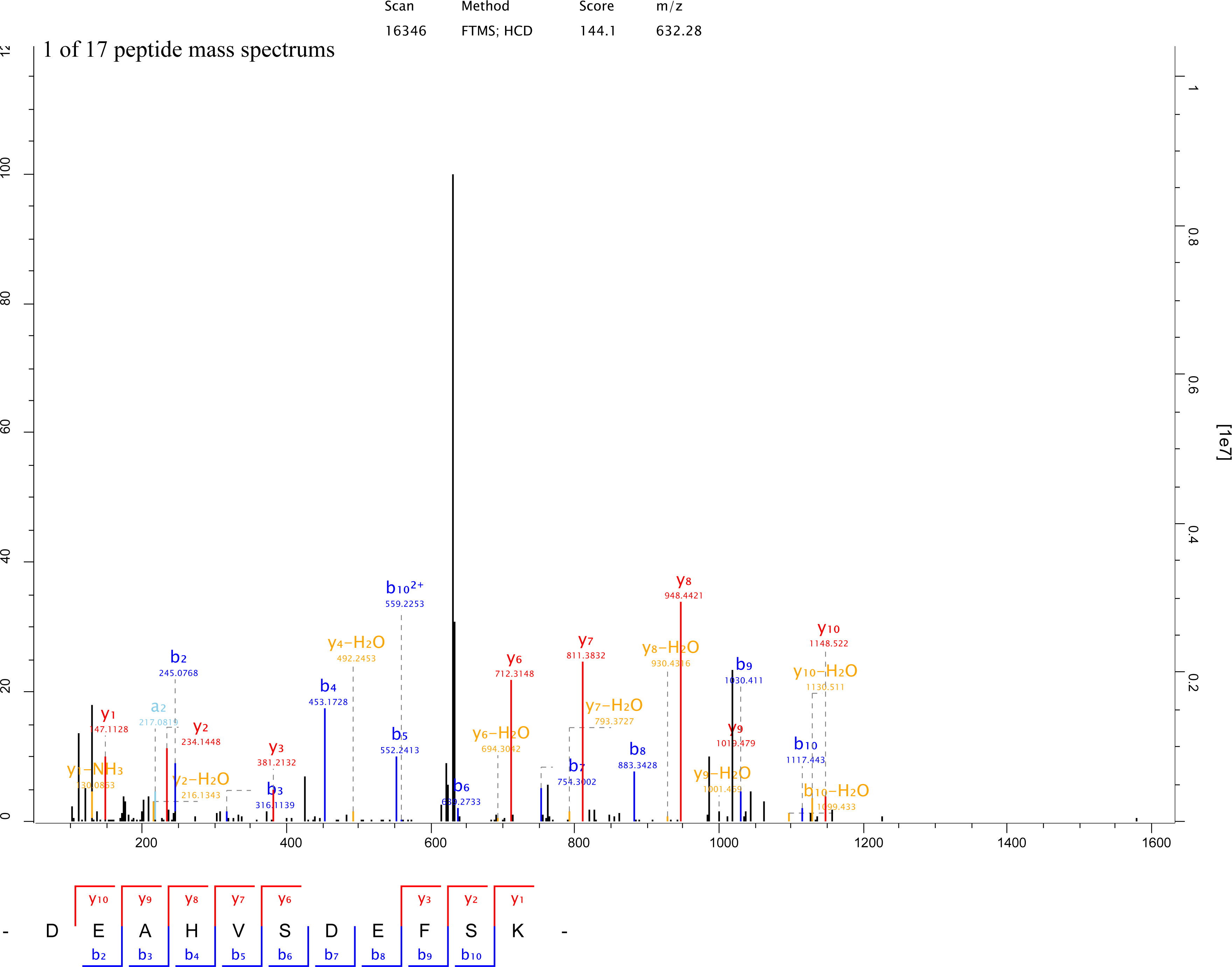
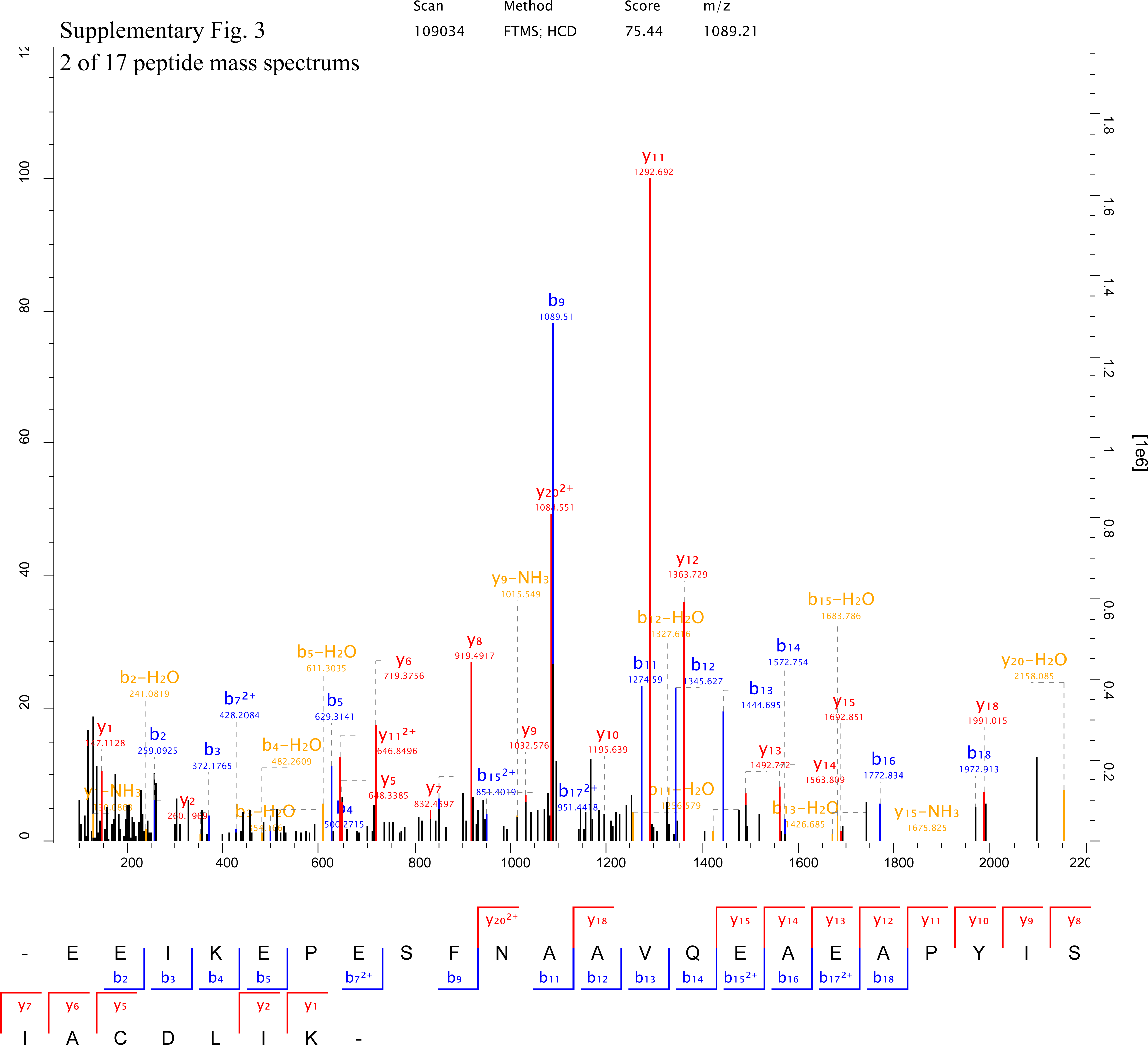
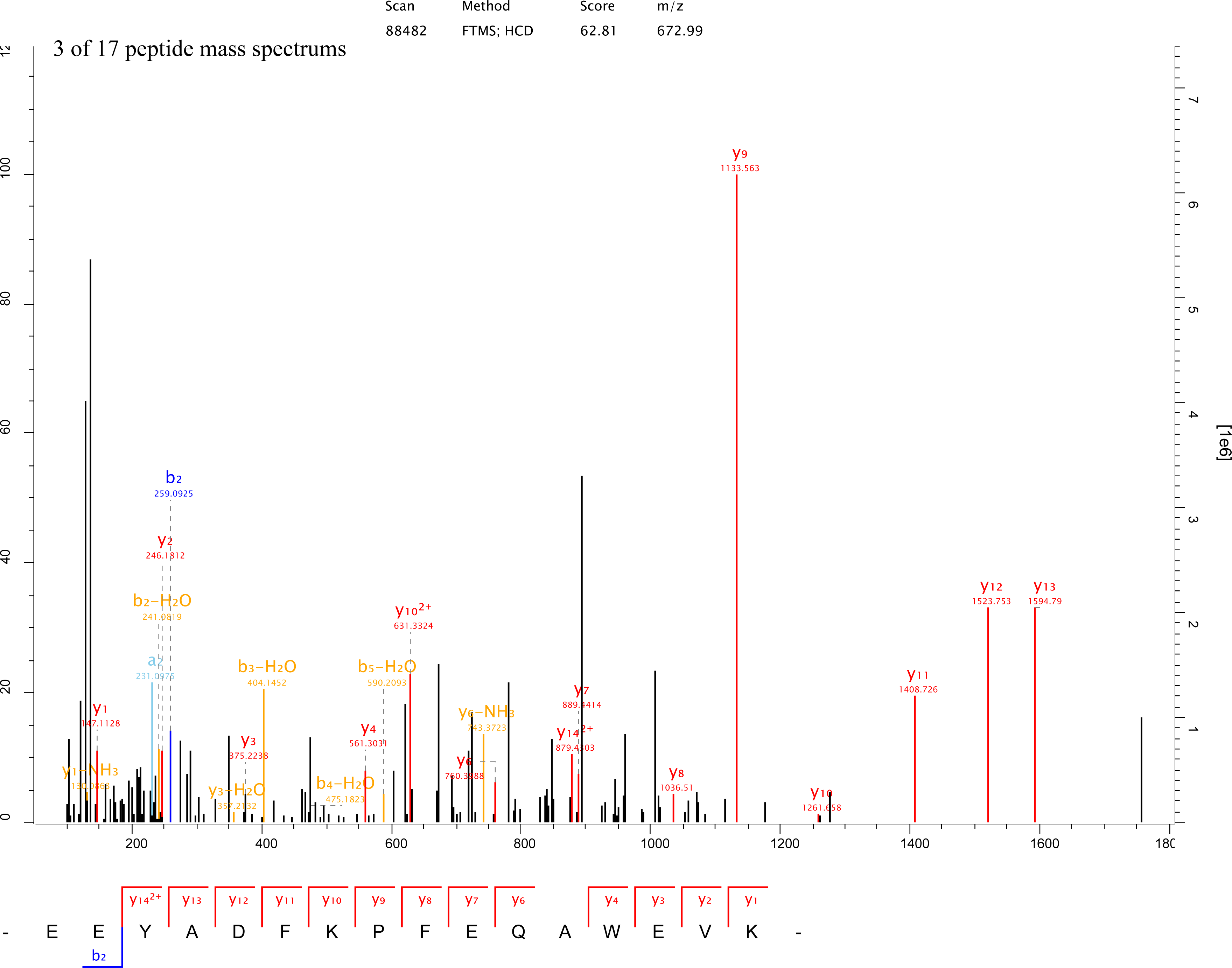
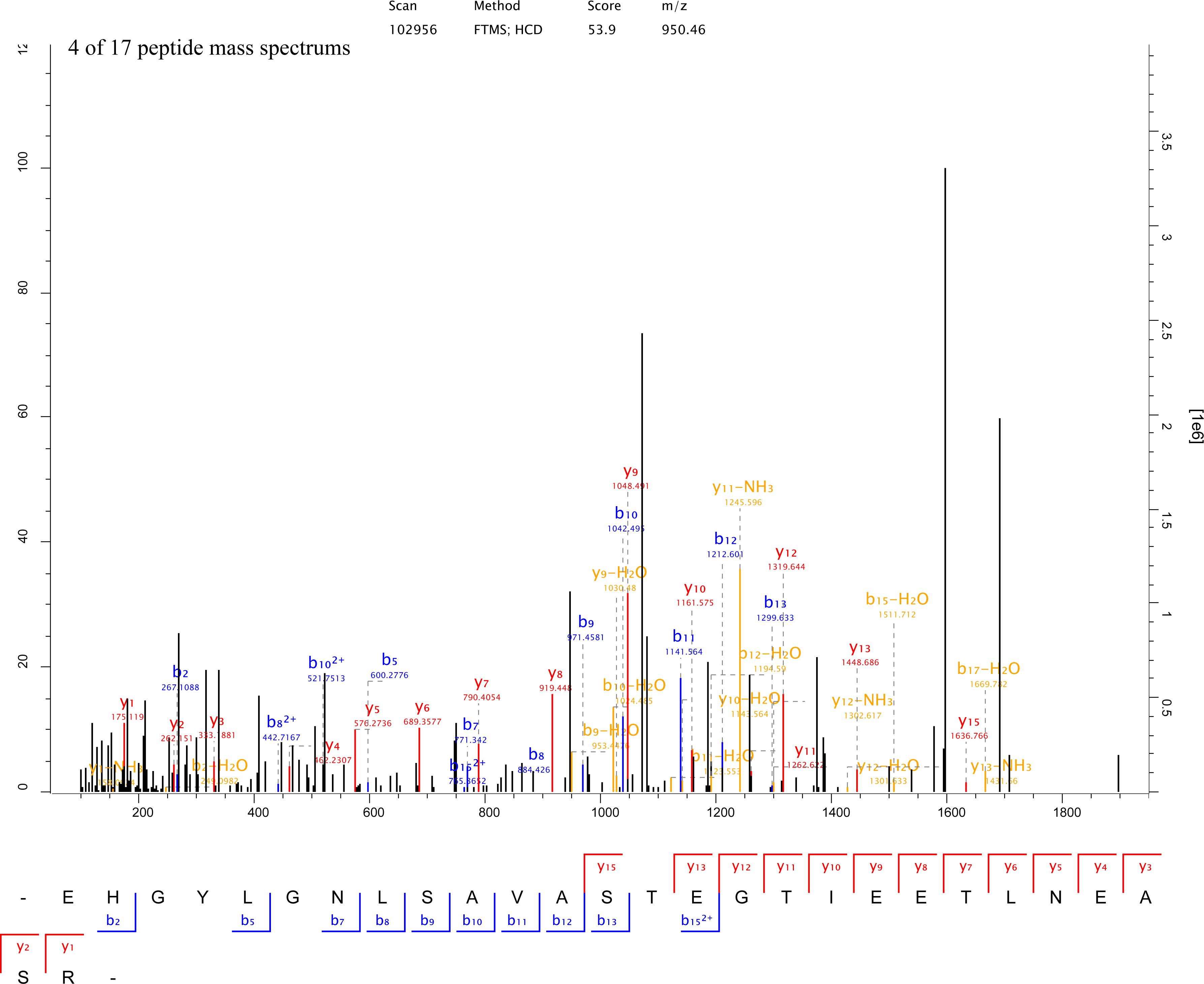
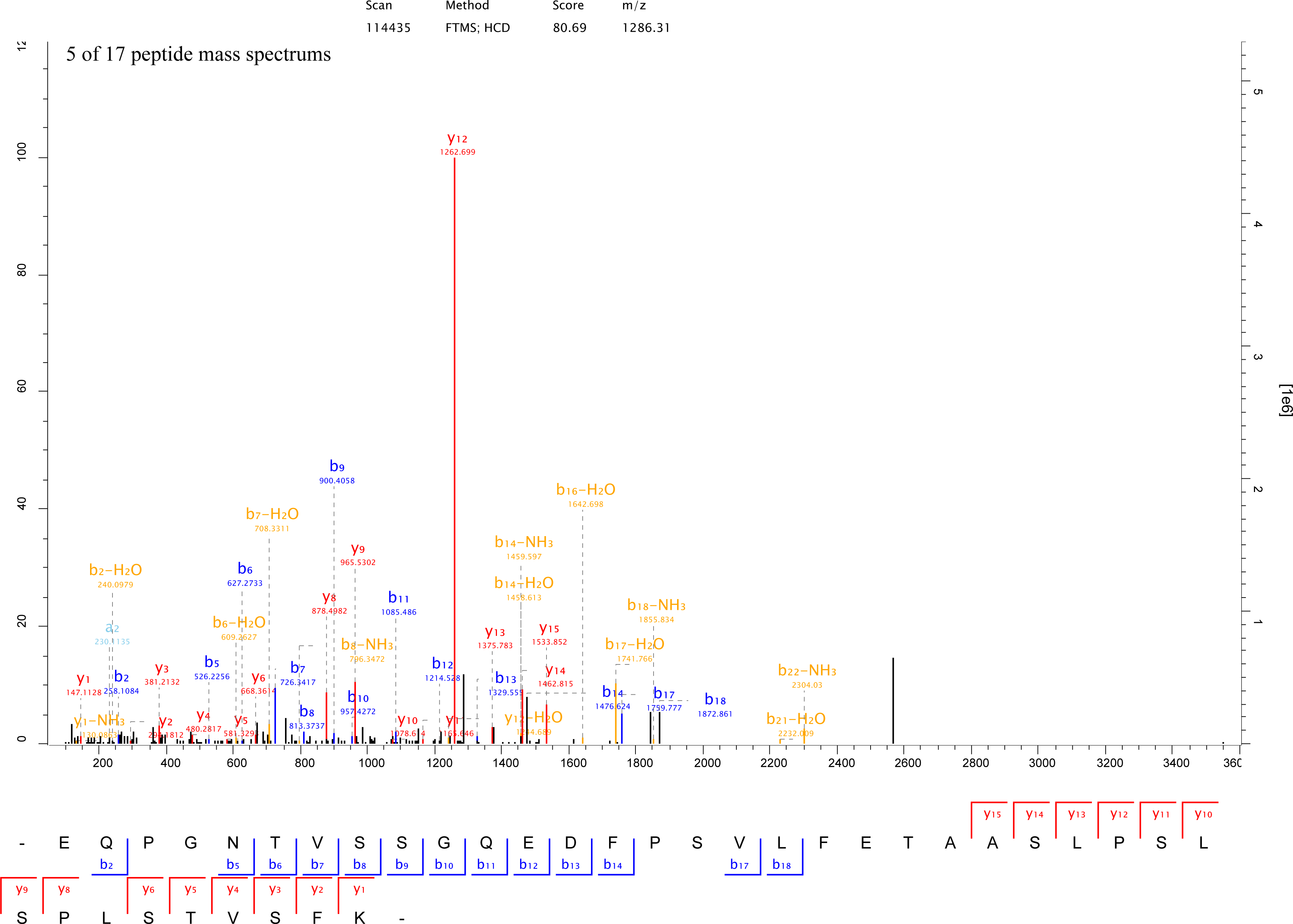
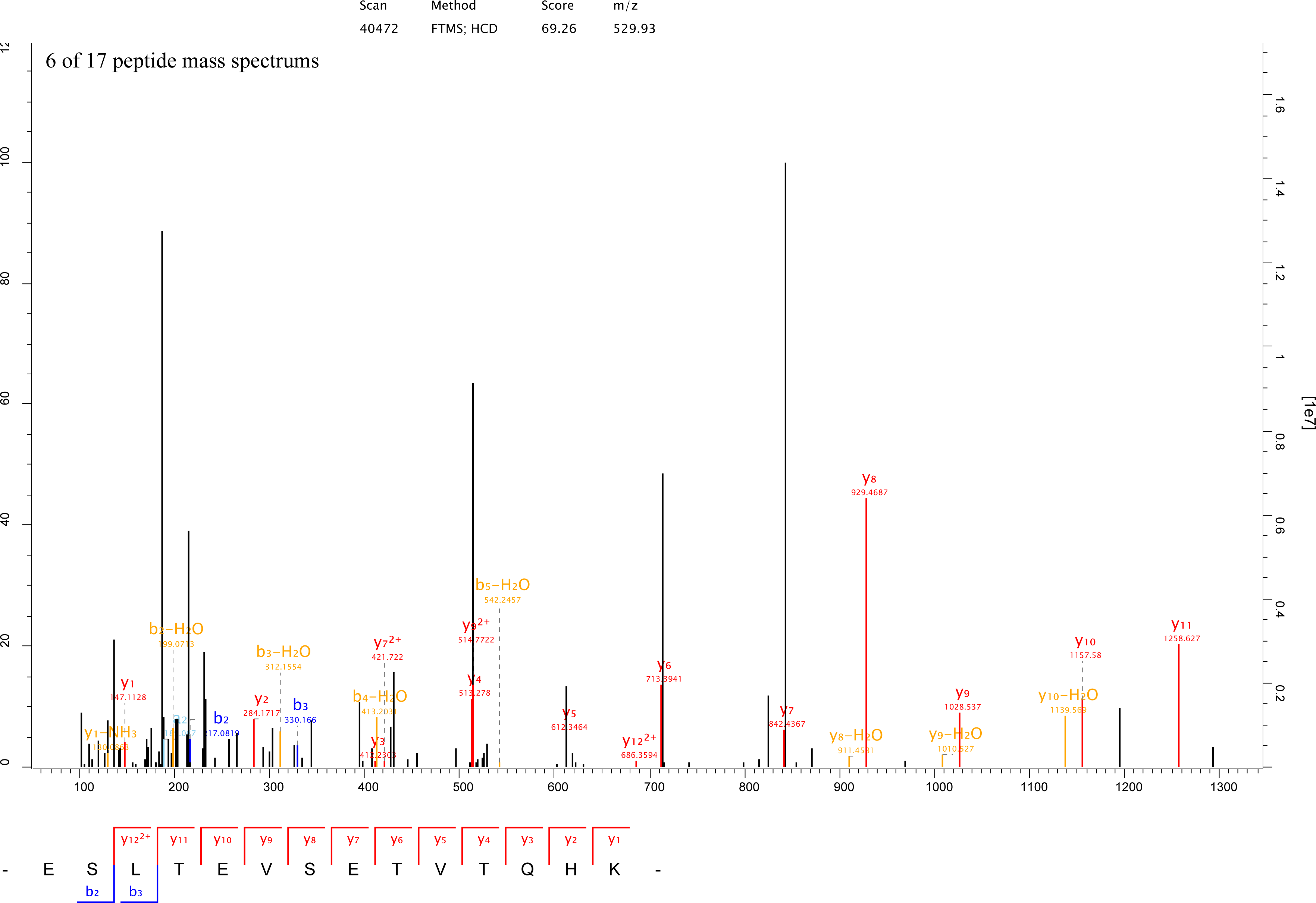
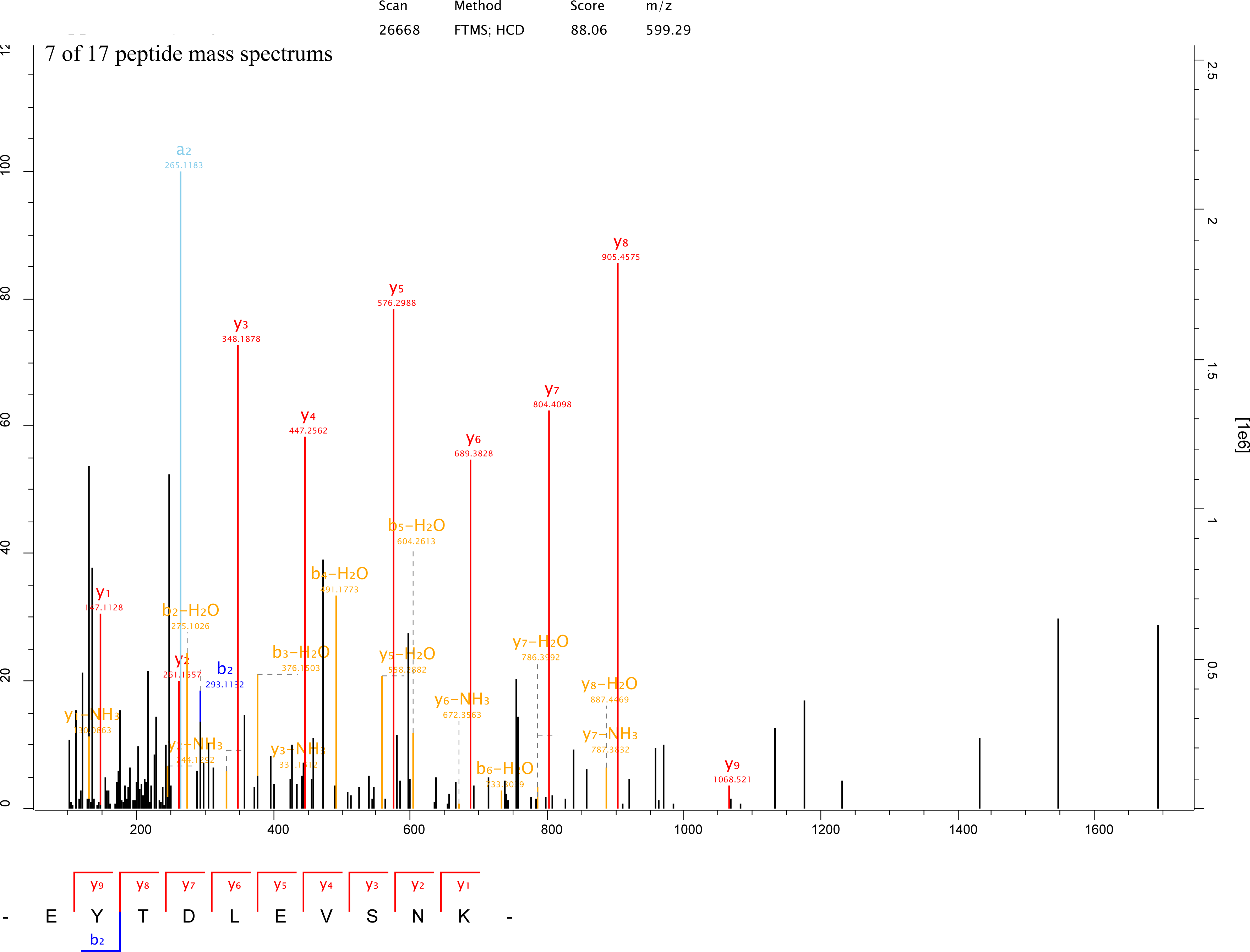
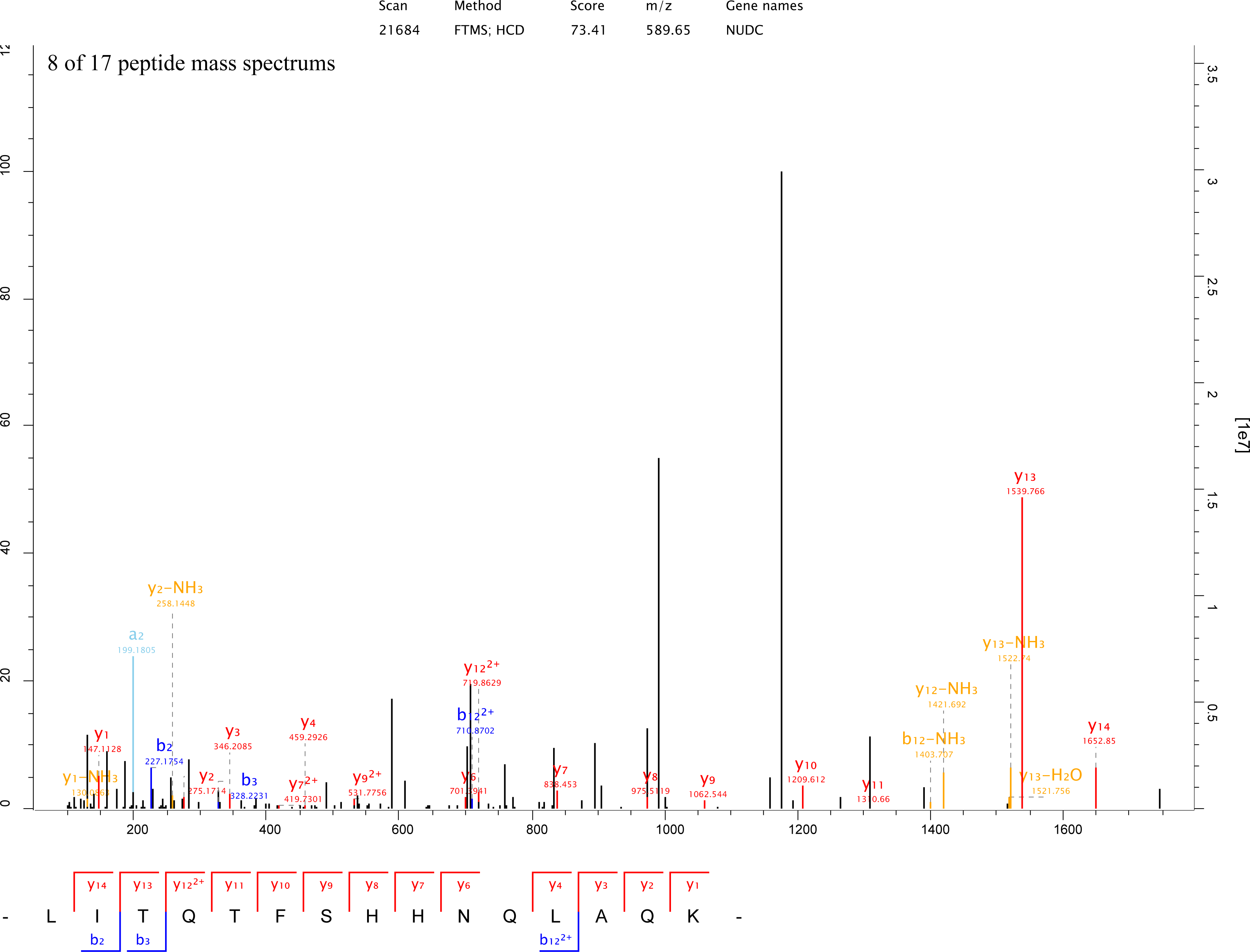
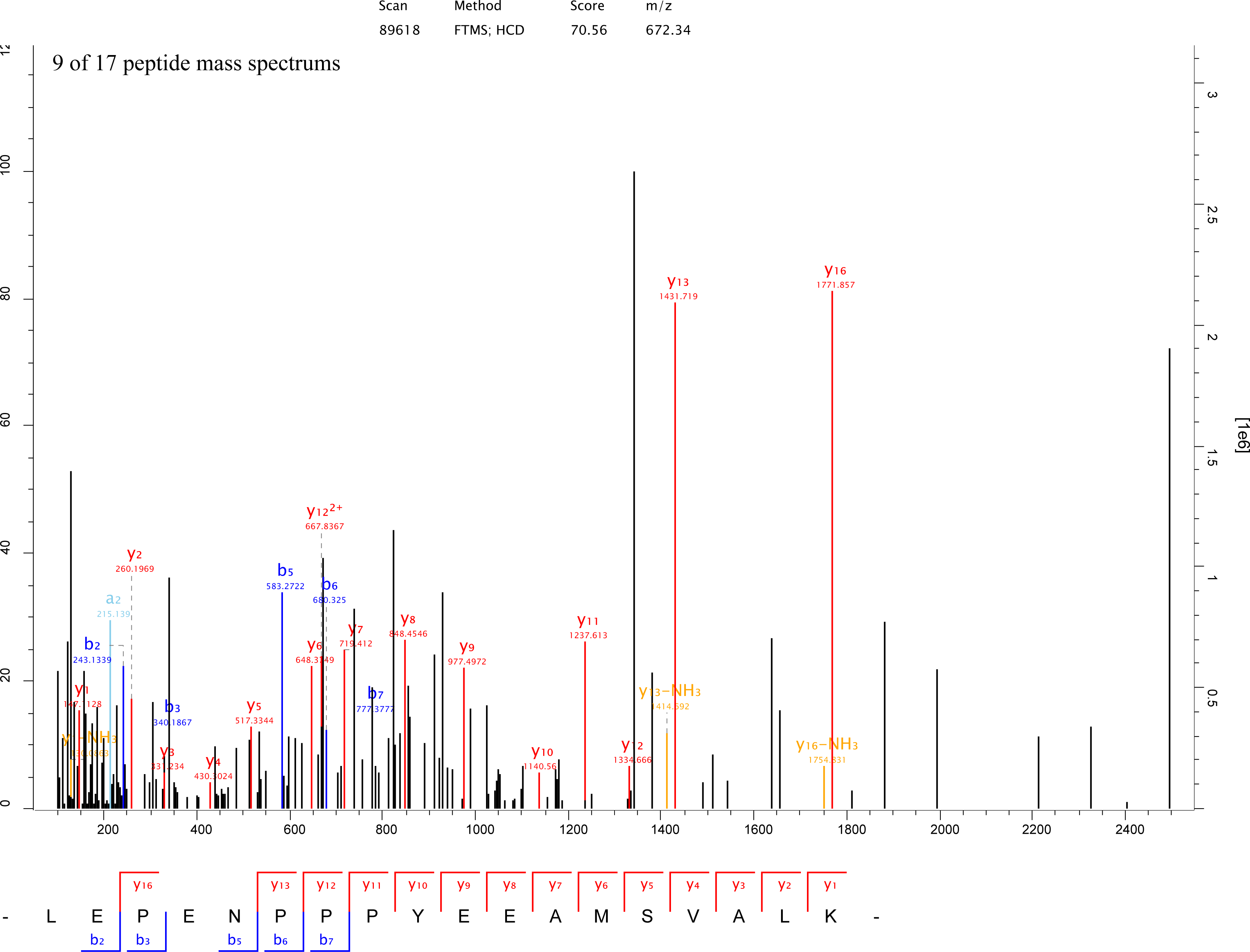
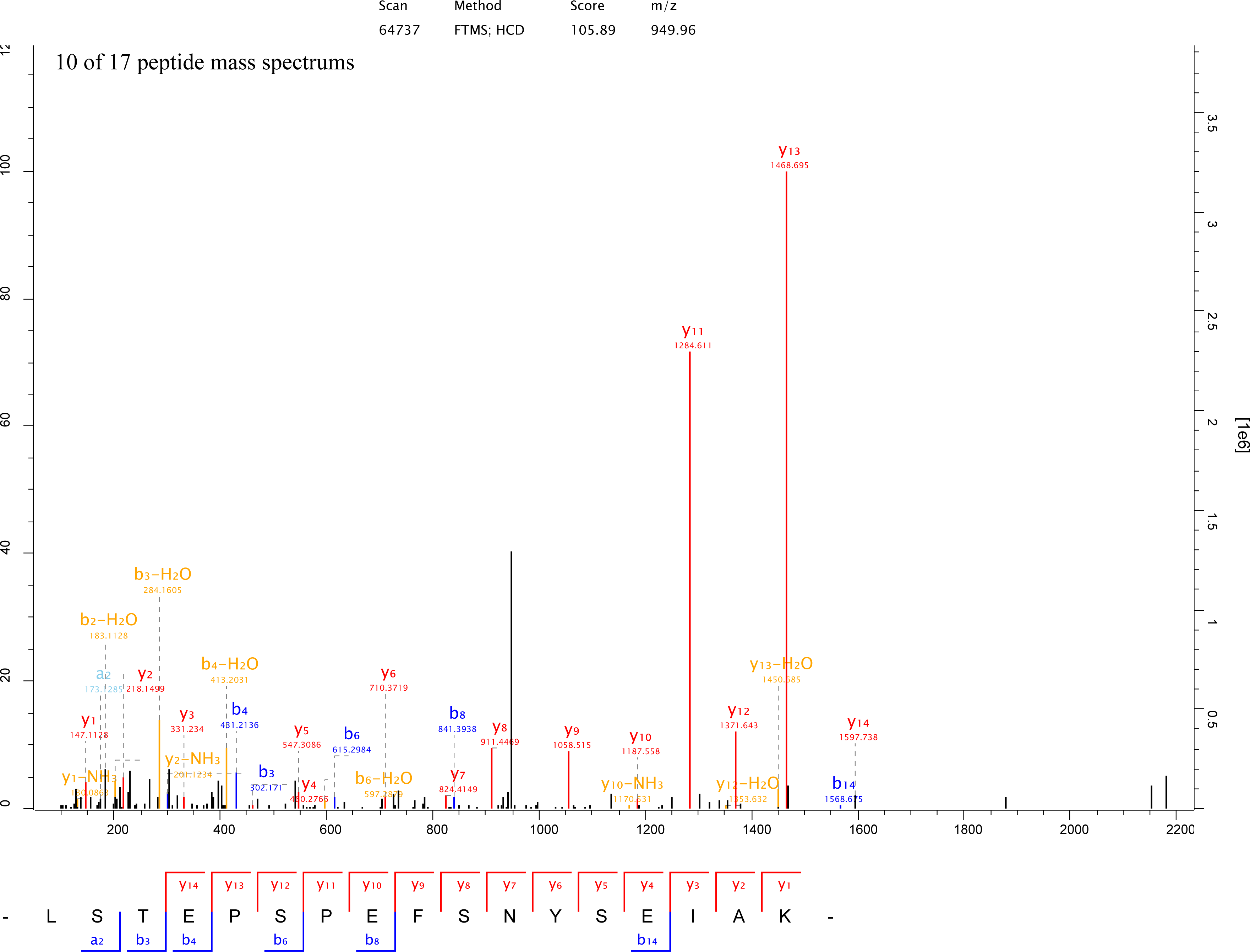
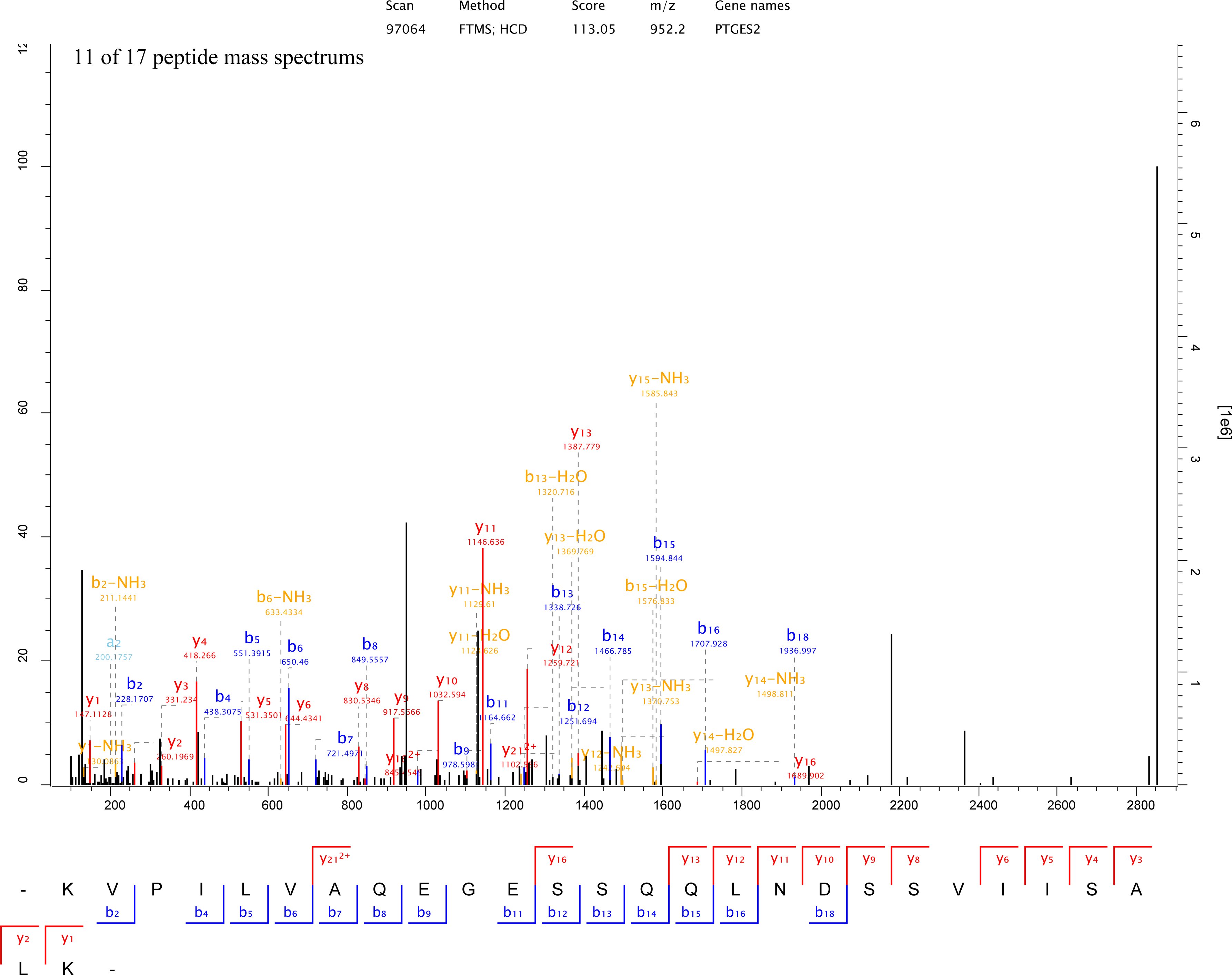
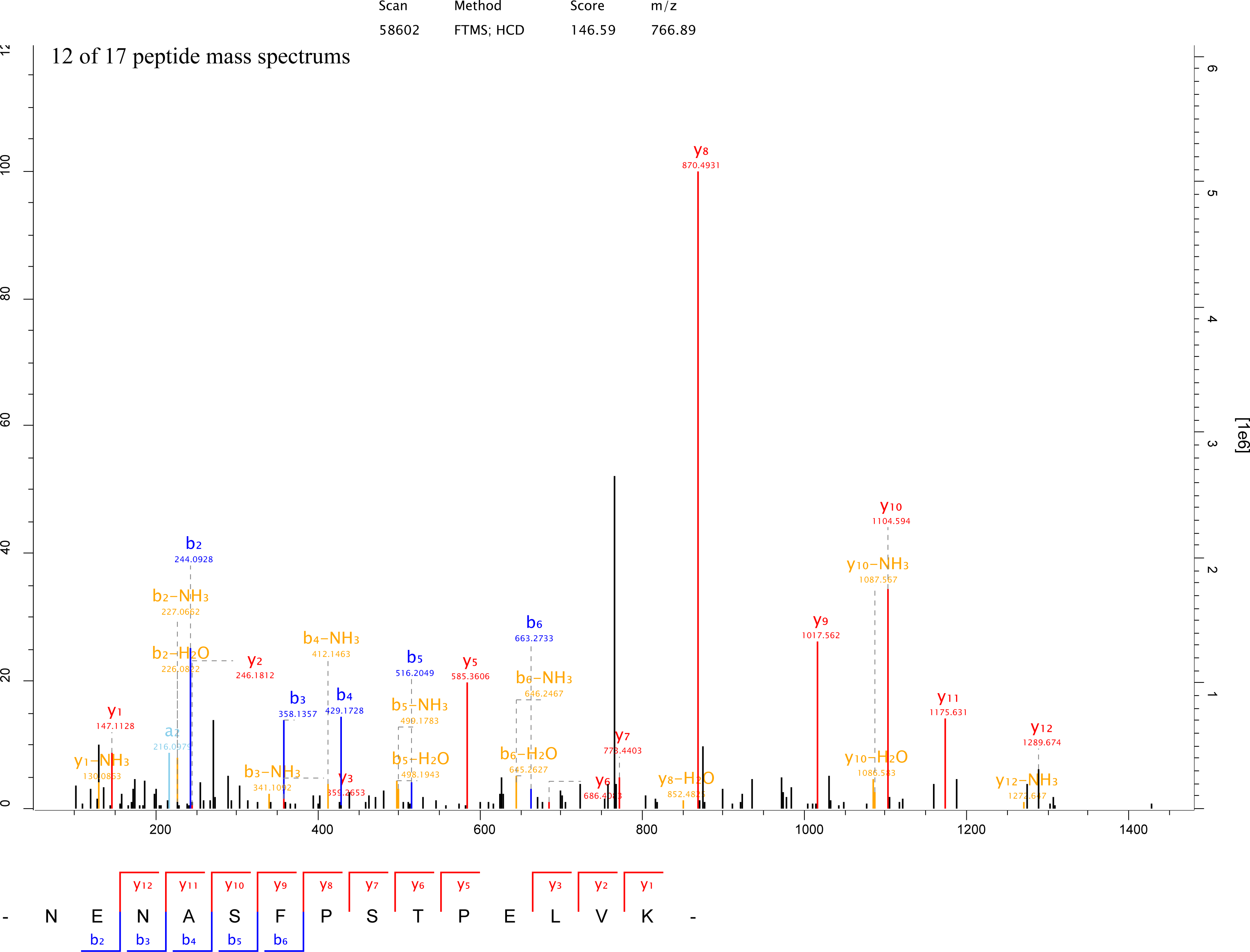
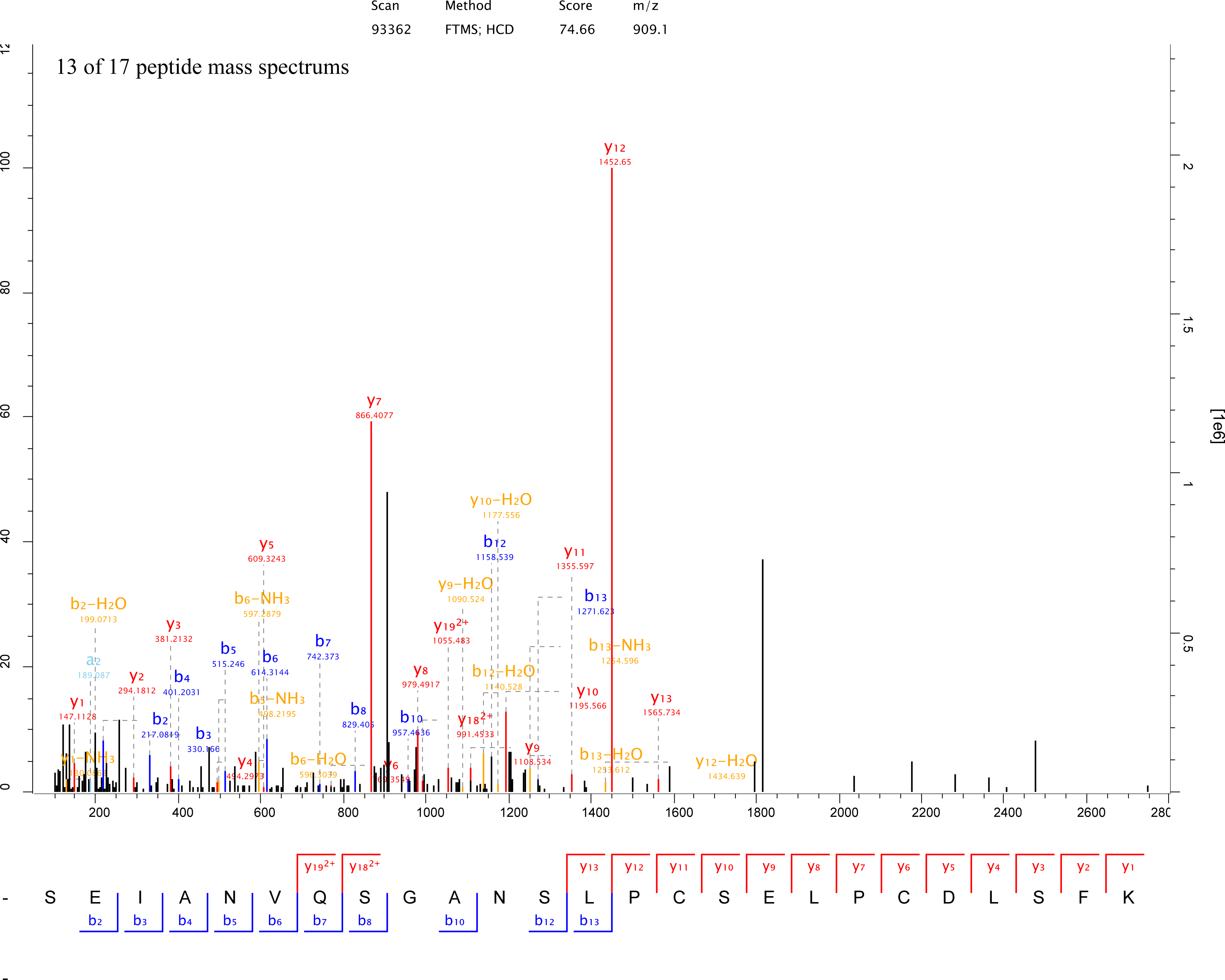
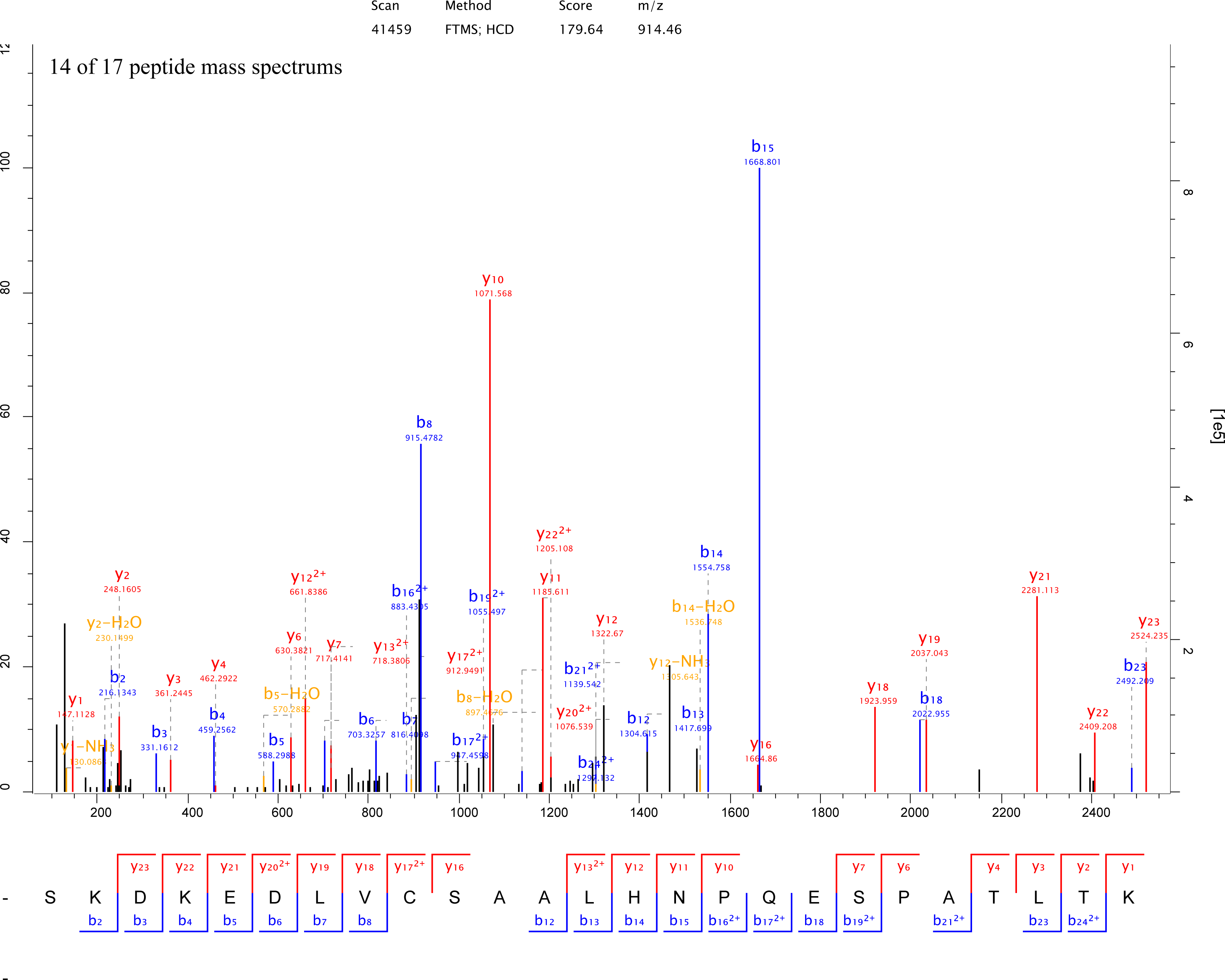
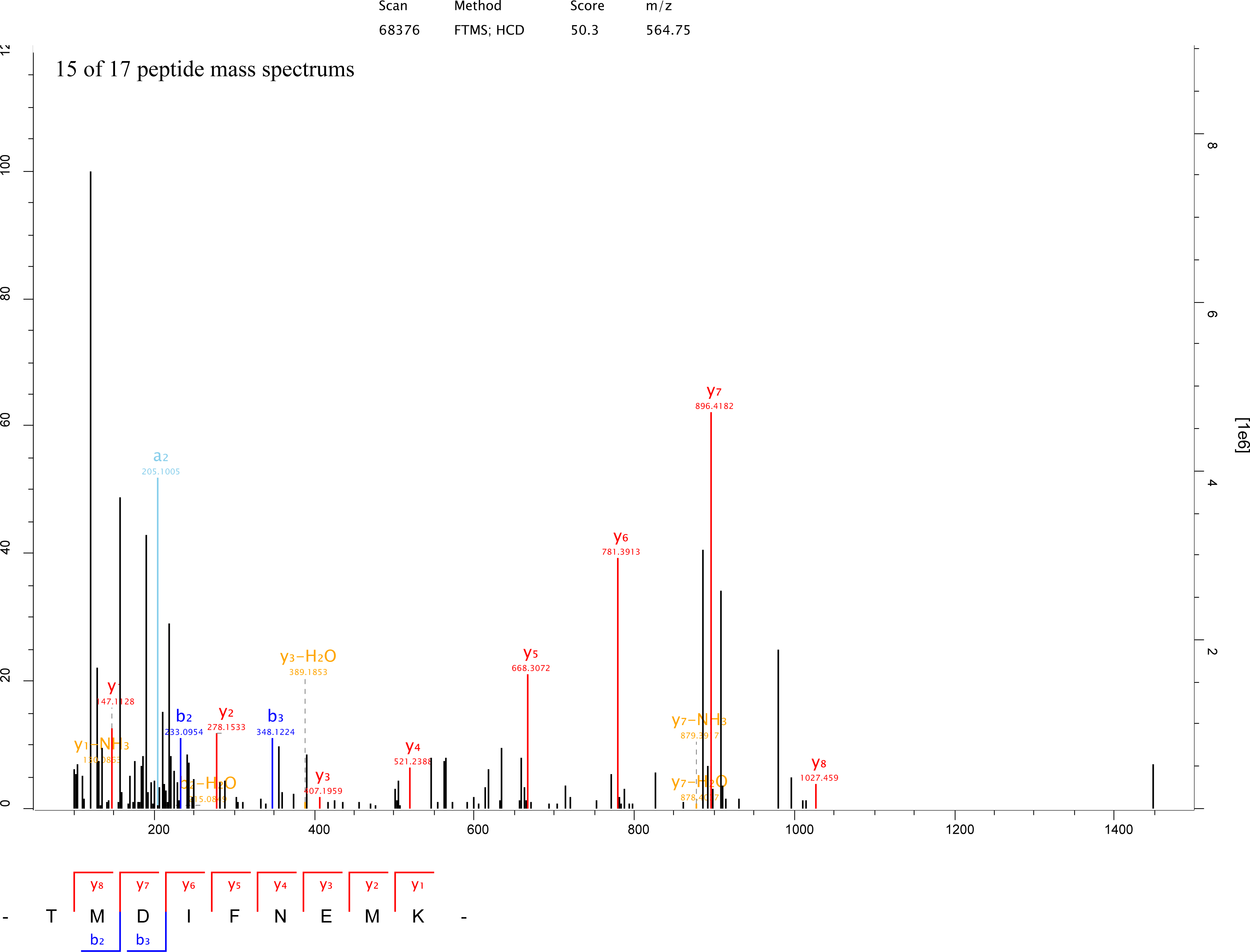
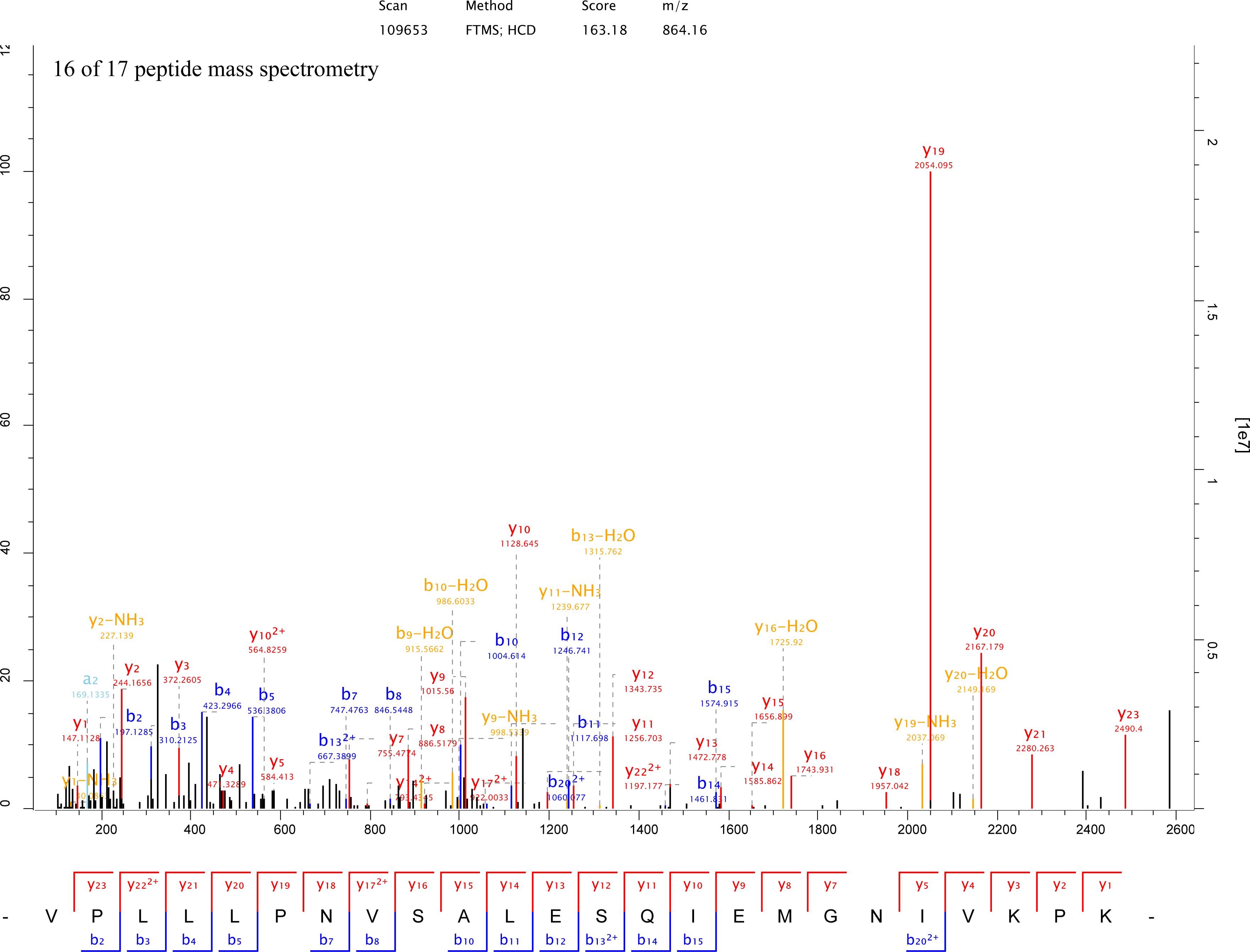
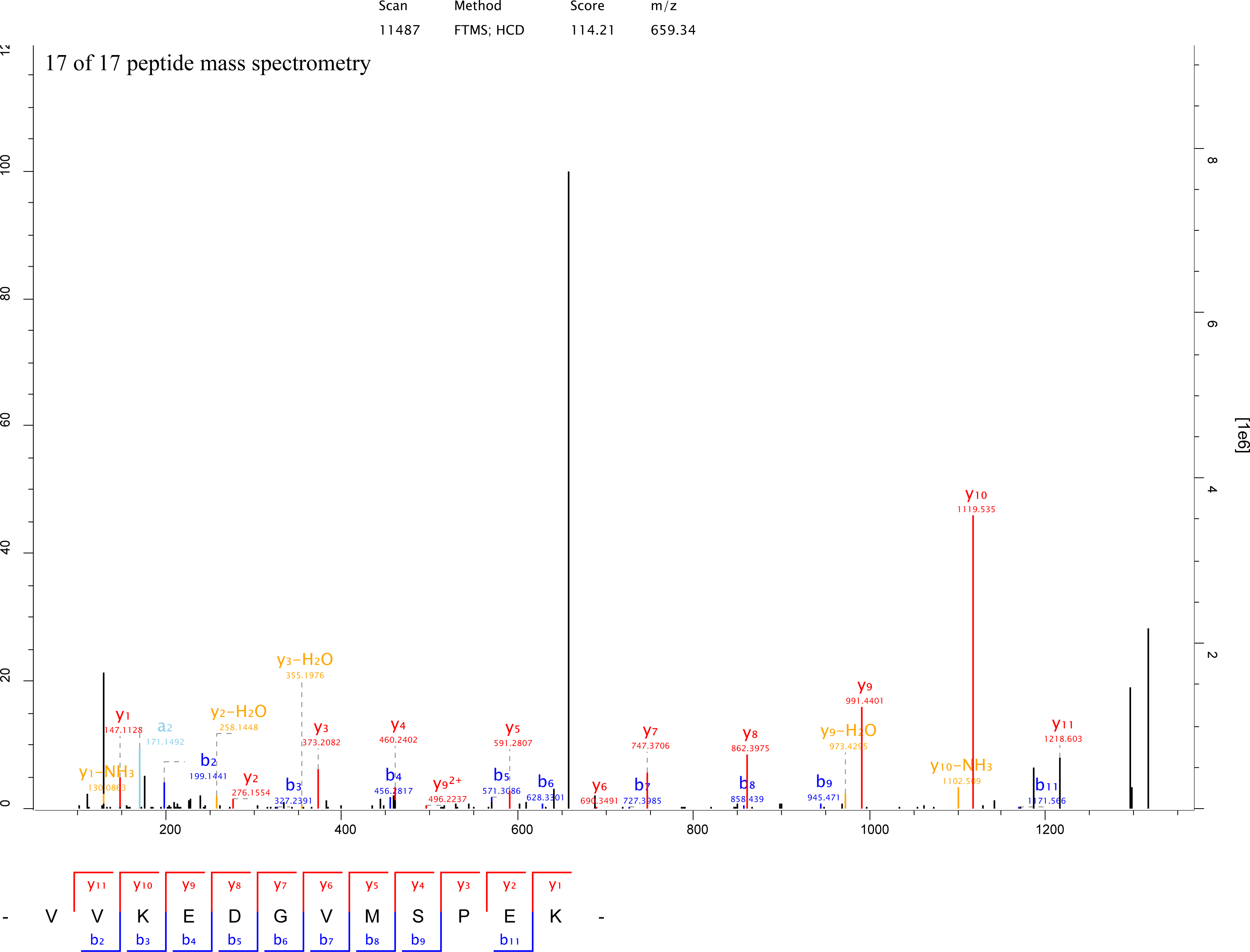
17 mass spectrums of the tryptic peptides of Rtn4 circRNA derived protein Left ordinate is the relative intensity, right ordinate is the absolute intensity, Horizontal ordinate is m/z.

**Supplementary Fig. 4.**
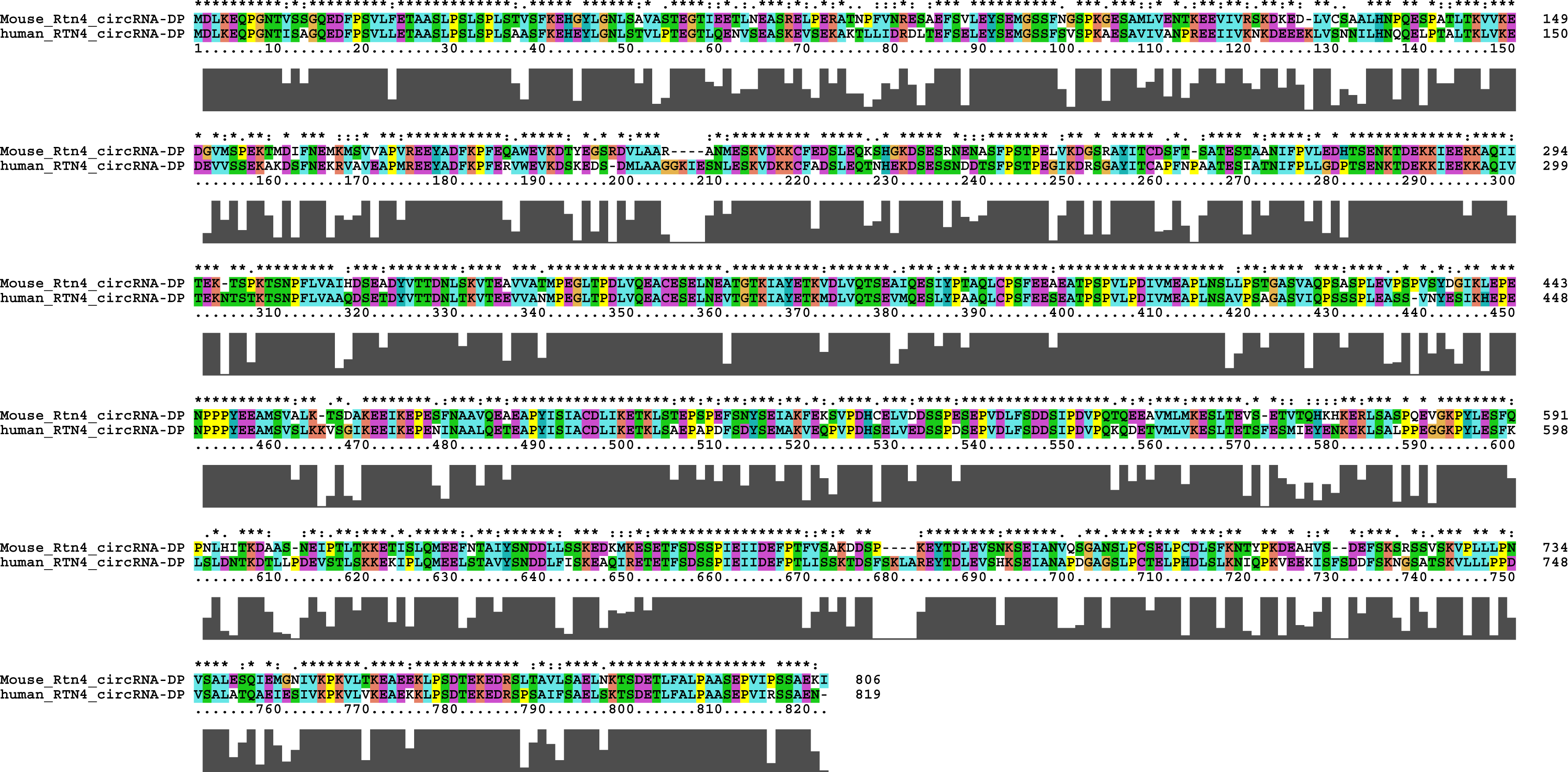
Alignment of RTN4 circRNA derived protein sequences from mouse and human. The sequences of mouse Rtn4 circRNA derived protein (Mouse_Rtn4_circRNA-DP) and human Rtn4 circRNA derived protein (human_Rtn4_circRNA-DP) were aligned through ClustalX2.

**Supplementary Fig. 5.**
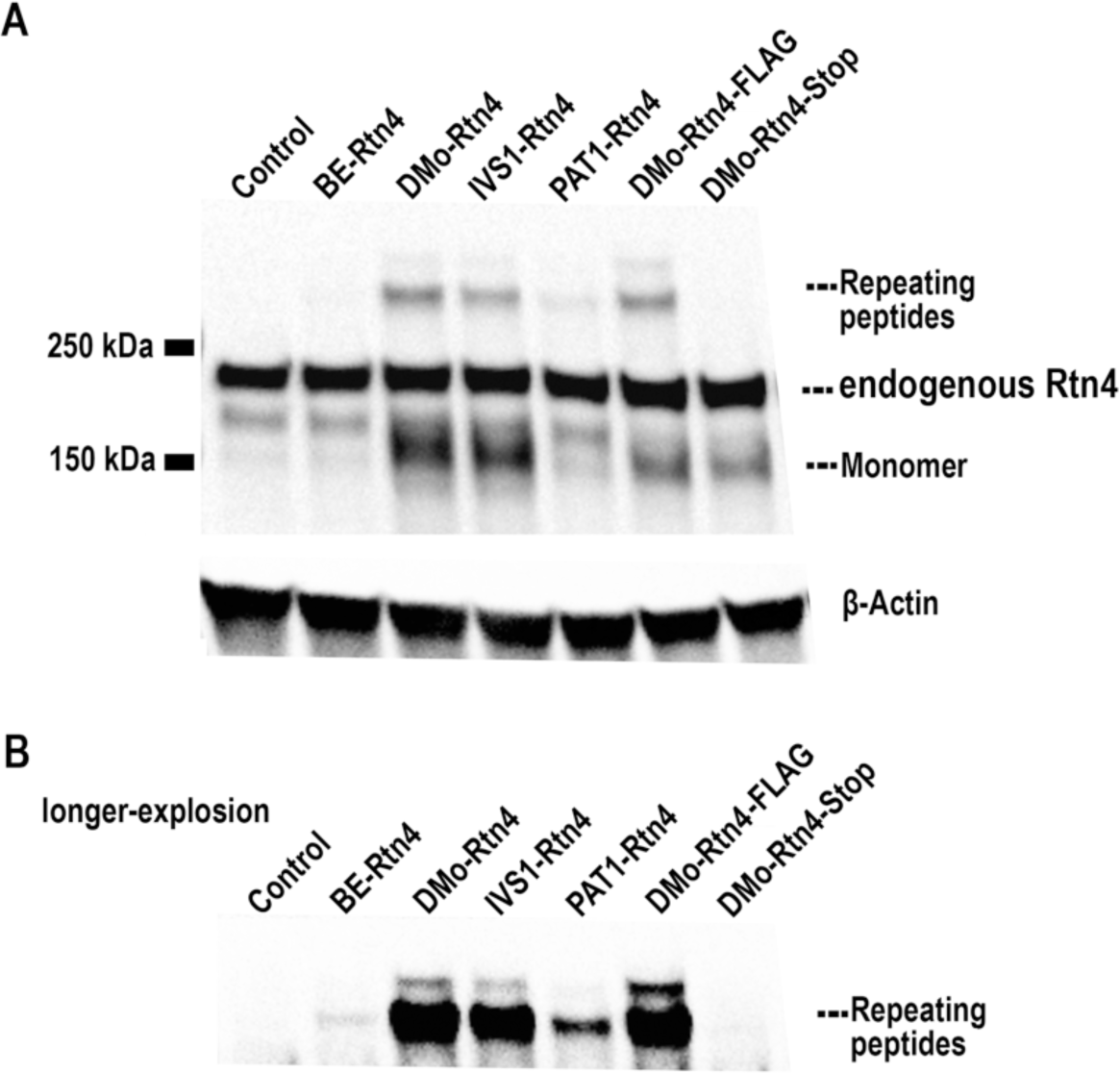
Rtn4 circRNA translation in N2a cells. A. Western blot with Anti-Nogo A antibody of N2a cells with Rtn4 circRNA overexpression. Control, empty vector; BE-Rtn4, pCircRNA-BE-Rtn4; DMo-Rtn4, pCircRNA-DMo-Rtn4; IVS1-Rtn4, pCircRNA-IVS1-Rtn4; PAT1-Rtn4, pCircRNA-PAT1-Rtn4; DMo-Rtn4-FLAG, pCircRNA-DMo-Rtn4-FLAG; DMo-Rtn4-Stop, pCircRNA-DMo-Rtn4-Stop; the high molecular weight bands bigger than 250 kDa represent the repeating peptides from Rtn4 circRNA continuous translation; monomer is the single round Rtn4 circRNA translation product; actin is used as loading control. B. the longer explosion of the high molecular weight bands bigger than 250 kDa.

## References

1. Chen, L.L. (2016) The biogenesis and emerging roles of circular RNAs. Nat Rev Mol Cell Biol, 17, 205–211.

2. Hentze, M.W. and Preiss, T. (2013) Circular RNAs: splicing’s enigma variations. EMBO J, 32, 923–925.

3. Vicens, Q. and Westhof, E. (2014) Biogenesis of Circular RNAs. Cell, 159, 13–14.

4. Rybak-Wolf, A., Stottmeister, C., Glazar, P., Jens, M., Pino, N., Giusti, S., Hanan, M., Behm, M., Bartok, O., Ashwal-Fluss, R. et al. (2015) Circular RNAs in the Mammalian Brain Are Highly Abundant, Conserved, and Dynamically Expressed. Mol Cell, 58, 870–885.

5. Veno, M.T., Hansen, T.B., Veno, S.T., Clausen, B.H., Grebing, M., Finsen, B., Holm, I.E. and Kjems, J. (2015) Spatio-temporal regulation of circular RNA expression during porcine embryonic brain development. Genome Biol, 16, 245.

6. You, X., Vlatkovic, I., Babic, A., Will, T., Epstein, I., Tushev, G., Akbalik, G., Wang, M., Glock, C., Quedenau, C. et al. (2015) Neural circular RNAs are derived from synaptic genes and regulated by development and plasticity. Nat Neurosci, 18, 603–610.

7. Westholm, J.O., Miura, P., Olson, S., Shenker, S., Joseph, B., Sanfilippo, P., Celniker, S.E., Graveley, B.R. and Lai, E.C. (2014) Genome-wide analysis of drosophila circular RNAs reveals their structural and sequence properties and age-dependent neural accumulation. Cell Rep, 9, 1966–1980.

8. Gruner, H., Cortes-Lopez, M., Cooper, D.A., Bauer, M. and Miura, P. (2016) CircRNA accumulation in the aging mouse brain. Sci Rep, 6, 38907.

9. Preusser, C., Hung, L.H., Schneider, T., Schreiner, S., Hardt, M., Moebus, A., Santoso, S. and Bindereif, A. (2018) Selective release of circRNAs in platelet-derived extracellular vesicles. Journal of extracellular vesicles, 7, 1424473.

10. Szabo, L., Morey, R., Palpant, N.J., Wang, P.L., Afari, N., Jiang, C., Parast, M.M., Murry, C.E., Laurent, L.C. and Salzman, J. (2015) Statistically based splicing detection reveals neural enrichment and tissue-specific induction of circular RNA during human fetal development. Genome Biol, 16, 126.

11. Salzman, J., Chen, R.E., Olsen, M.N., Wang, P.L. and Brown, P.O. (2013) Cell-type specific features of circular RNA expression. PLoS genetics, 9, e1003777.

12. Conn, S.J., Pillman, K.A., Toubia, J., Conn, V.M., Salmanidis, M., Phillips, C.A., Roslan, S., Schreiber, A.W., Gregory, P.A. and Goodall, G.J. (2015) The RNA binding protein quaking regulates formation of circRNAs. Cell, 160, 1125–1134.

13. Hansen, T.B., Jensen, T.I., Clausen, B.H., Bramsen, J.B., Finsen, B., Damgaard, C.K. and Kjems, J. (2013) Natural RNA circles function as efficient microRNA sponges. Nature, 495, 384–388.

14. Memczak, S., Jens, M., Elefsinioti, A., Torti, F., Krueger, J., Rybak, A., Maier, L., Mackowiak, S.D., Gregersen, L.H., Munschauer, M. et al. (2013) Circular RNAs are a large class of animal RNAs with regulatory potency. Nature, 495, 333–338.

15. Piwecka, M., Glazar, P., Hernandez-Miranda, L.R., Memczak, S., Wolf, S.A., Rybak-Wolf, A., Filipchyk, A., Klironomos, F., Cerda Jara, C.A., Fenske, P. et al. (2017) Loss of a mammalian circular RNA locus causes miRNA deregulation and affects brain function. Science.

16. Kristensen, L.S., Hansen, T.B., Veno, M.T. and Kjems, J. (2017) Circular RNAs in cancer: opportunities and challenges in the field. Oncogene.

17. Guarnerio, J., Bezzi, M., Jeong, J.C., Paffenholz, S.V., Berry, K., Naldini, M.M., Lo-Coco, F., Tay, Y., Beck, A.H. and Pandolfi, P.P. (2016) Oncogenic Role of Fusion-circRNAs Derived from Cancer-Associated Chromosomal Translocations. Cell, 165, 289–302.

18. Xia, S., Feng, J., Chen, K., Ma, Y., Gong, J., Cai, F., Jin, Y., Gao, Y., Xia, L., Chang, H. et al. (2017) CSCD: a database for cancer-specific circular RNAs. Nucleic Acids Res.

19. Zhang, X.O., Wang, H.B., Zhang, Y., Lu, X., Chen, L.L. and Yang, L. (2014) Complementary sequence-mediated exon circularization. Cell, 159, 134–147.

20. Liang, D. and Wilusz, J.E. (2014) Short intronic repeat sequences facilitate circular RNA production. Genes Dev, 28, 2233–2247.

21. Ashwal-Fluss, R., Meyer, M., Pamudurti, N.R., Ivanov, A., Bartok, O., Hanan, M., Evantal, N., Memczak, S., Rajewsky, N. and Kadener, S. (2014) circRNA biogenesis competes with pre-mRNA splicing. Mol Cell, 56, 55–66.

22. Starke, S., Jost, I., Rossbach, O., Schneider, T., Schreiner, S., Hung, L.H. and Bindereif, A. (2015) Exon circularization requires canonical splice signals. Cell Rep, 10, 103–111.

23. Zhang, Y., Xue, W., Li, X., Zhang, J., Chen, S., Zhang, J.L., Yang, L. and Chen, L.L. (2016) The Biogenesis of Nascent Circular RNAs. Cell Rep, 15, 611–624.

24. Legnini, I. and Bozzoni, I. (2017), Essentials of Noncoding RNA in Neuroscience. Academic Press, pp. 247–263.

25. Buchman, A.R. and Berg, P. (1988) Comparison of intron-dependent and intron-independent gene expression. Mol Cell Biol, 8, 4395–4405.

26. Huang, M.T. and Gorman, C.M. (1990) Intervening sequences increase efficiency of RNA 3’ processing and accumulation of cytoplasmic RNA. Nucleic Acids Res, 18, 937–947.

27. Brinster, R.L., Allen, J.M., Behringer, R.R., Gelinas, R.E. and Palmiter, R.D. (1988) Introns increase transcriptional efficiency in transgenic mice. Proc Natl Acad Sci U S A, 85, 836–840.

28. Choi, T., Huang, M., Gorman, C. and Jaenisch, R. (1991) A generic intron increases gene expression in transgenic mice. Mol Cell Biol, 11, 3070–3074.

29. Palmiter, R.D., Sandgren, E.P., Avarbock, M.R., Allen, D.D. and Brinster, R.L. (1991) Heterologous introns can enhance expression of transgenes in mice. Proc Natl Acad Sci U S A, 88, 478–482.

30. Gallegos, J.E. and Rose, A.B. (2015) The enduring mystery of intron-mediated enhancement. Plant Sci, 237, 8–15.

31. Rose, A.B. and Beliakoff, J.A. (2000) Intron-mediated enhancement of gene expression independent of unique intron sequences and splicing. Plant Physiol, 122, 535–542.

32. Laxa, M. (2016) Intron-Mediated Enhancement: A Tool for Heterologous Gene Expression in Plants? Front Plant Sci, 7, 1977.

33. Schwab, M.E. (2010) Functions of Nogo proteins and their receptors in the nervous system. Nat Rev Neurosci, 11, 799–811.

34. Seiler, S., Di Santo, S. and Widmer, H.R. (2016) Non-canonical actions of Nogo-A and its receptors. Biochem Pharmacol, 100, 28–39.

35. Thinakaran, G., Teplow, D.B., Siman, R., Greenberg, B. and Sisodia, S.S. (1996) Metabolism of the “Swedish” amyloid precursor protein variant in neuro2a (N2a) cells. Evidence that cleavage at the “beta-secretase” site occurs in the golgi apparatus. J Biol Chem, 271, 9390–9397.

36. Livak, K.J. and Schmittgen, T.D. (2001) Analysis of relative gene expression data using real-time quantitative PCR and the 2(-Delta Delta C(T)) Method. Methods, 25, 402–408.

37. Schneider, T., Schreiner, S., Preusser, C., Bindereif, A. and Rossbach, O. (2018) Northern Blot Analysis of Circular RNAs. Methods Mol Biol, 1724, 119–133.

38. Li, X. and Franz, T. (2014) Up to date sample preparation of proteins for mass spectrometric analysis. Arch Physiol Biochem, 120, 188–191.

39. Kulak, N.A., Pichler, G., Paron, I., Nagaraj, N. and Mann, M. (2014) Minimal, encapsulated proteomic-sample processing applied to copy-number estimation in eukaryotic cells. Nat Methods, 11, 319–324.

40. Cox, J., Neuhauser, N., Michalski, A., Scheltema, R.A., Olsen, J.V. and Mann, M. (2011) Andromeda: a peptide search engine integrated into the MaxQuant environment. J Proteome Res, 10, 1794–1805.

41. Graille, M. and Seraphin, B. (2012) Surveillance pathways rescuing eukaryotic ribosomes lost in translation. Nat Rev Mol Cell Biol, 13, 727–735.

42. Lykke-Andersen, S. and Jensen, T.H. (2015) Nonsense-mediated mRNA decay: an intricate machinery that shapes transcriptomes. Nat Rev Mol Cell Biol, 16, 665–677.

43. Guan, Y., Zhu, Q., Huang, D., Zhao, S., Jan Lo, L. and Peng, J. (2015) An equation to estimate the difference between theoretically predicted and SDS PAGE-displayed molecular weights for an acidic peptide. Sci Rep, 5, 13370.

44. Chen, C.Y. and Sarnow, P. (1995) Initiation of protein synthesis by the eukaryotic translational apparatus on circular RNAs. Science, 268, 415–417.

45. Abe, N., Matsumoto, K., Nishihara, M., Nakano, Y., Shibata, A., Maruyama, H., Shuto, S., Matsuda, A., Yoshida, M., Ito, Y. et al. (2015) Rolling Circle Translation of Circular RNA in Living Human Cells. Sci Rep, 5, 16435.

46. Wang, Y. and Wang, Z. (2015) Efficient backsplicing produces translatable circular mRNAs. RNA, 21, 172–179.

47. Legnini, I., Di Timoteo, G., Rossi, F., Morlando, M., Briganti, F., Sthandier, O., Fatica, A., Santini, T., Andronache, A., Wade, M. et al. (2017) Circ-ZNF609 Is a Circular RNA that Can Be Translated and Functions in Myogenesis. Mol Cell, 66, 22–37 e29.

48. Pamudurti, N.R., Bartok, O., Jens, M., Ashwal-Fluss, R., Stottmeister, C., Ruhe, L., Hanan, M., Wyler, E., Perez-Hernandez, D., Ramberger, E. et al. (2017) Translation of CircRNAs. Mol Cell, 66, 9–21 e27.

49. Yang, Y., Fan, X., Mao, M., Song, X., Wu, P., Zhang, Y., Jin, Y., Yang, Y., Chen, L.L., Wang, Y. et al. (2017) Extensive translation of circular RNAs driven by N6-methyladenosine. Cell Res, 27, 626–641.

50. Tatomer, D.C. and Wilusz, J.E. (2017) An Unchartered Journey for Ribosomes: Circumnavigating Circular RNAs to Produce Proteins. Mol Cell, 66, 1–2.

51. Schneider, T. and Bindereif, A. (2017) Circular RNAs: Coding or noncoding? Cell Res, 27, 724–725.

52. Jeck, W.R. and Sharpless, N.E. (2014) Detecting and characterizing circular RNAs. Nat Biotechnol, 32, 453–461.

53. Schwab, M.E. and Strittmatter, S.M. (2014) Nogo limits neural plasticity and recovery from injury. Curr Opin Neurobiol, 27, 53–60.

## References

1. Li, X. and Franz, T. (2014) Up to date sample preparation of proteins for mass spectrometric analysis. Arch Physiol Biochem, 120, 188–191.

2. Kulak, N.A., Pichler, G., Paron, I., Nagaraj, N. and Mann, M. (2014) Minimal, encapsulated proteomic-sample processing applied to copy-number estimation in eukaryotic cells. Nat Methods, 11, 319–324.

3. Cox, J., Neuhauser, N., Michalski, A., Scheltema, R.A., Olsen, J.V. and Mann, M. (2011) Andromeda: a peptide search engine integrated into the MaxQuant environment. J Proteome Res, 10, 1794–1805.

